# Multi-sensory integration in the mouse cortical connectome using a network diffusion model

**DOI:** 10.1101/832485

**Authors:** Kamal Shadi, Eva Dyer, Constantine Dovrolis

## Abstract

Having a structural network representation of connectivity in the brain is instrumental in analyzing communication dynamics and information processing in the brain. In this work, we make steps towards understanding multi-sensory information flow and integration using a network diffusion approach. In particular, we model the flow of evoked activity, initiated by stimuli at primary sensory regions, using the *Asynchronous Linear Threshold (ALT) diffusion model*. The ALT model captures how evoked activity that originates at a given region of the cortex “ripples through” other brain regions (referred to as an *activation cascade*). By comparing the model results to functional datasets based on Voltage Sensitive Dye (VSD) imaging, we find that in most cases the ALT model predicts the temporal ordering of an activation cascade correctly. Our results on the Mouse Connectivity Atlas from the Allen Institute for Brain Science show that a small number of brain regions are involved in many primary sensory streams – the claustrum and the parietal temporal cortex being at the top of the list. This suggests that the cortex relies on an *hourglass architecture* to first integrate and compress multi-sensory information from multiple sensory regions, before utilizing that lower-dimensionality representation in higher-level association regions and more complex cognitive tasks.

## 1 Introduction

Perception requires the integration of multiple sensory inputs across distributed areas throughout the brain [57]. While sensory integration at the behavioral level has been extensively studied [53], the network and system-level mechanisms underlying *Multi-Sensory Integration (MSI)* are still not well understood, especially in terms of the role that cortex plays. The traditional view of primary sensory areas as processing a single modality is rapidly shifting towards a view of the cortex as highly integrated and multi-sensory [32]. The somatosensory, visual, auditory, gustatory and other sensory streams, come together (integrate) and separate (diverge) to be processed in different parts of the cortex. The neural basis of how these sensory streams are processed and how they generate a coherent perceptual state remains elusive [55]. It is likely that this state is created and regulated by multiple structures distributed throughout the cortex working together in concert [13].

To understand the architectural principles that enable MSI, we need data and models that span the entire brain — focusing not on individual neurons, regions or even circuits, but on distributed networks. The connectome is thus a potentially powerful tool for studying MSI. However, it would not be enough to just know how different brain regions are connected anatomically. Rather, we need models that combine structure (connectomics) with function [2] to address the question of which connections and paths are “activated” by different sensory modalities. Having both a structural network and a functional model in hand, we can begin to tackle the problem of discovering the networks that support and constrain MSI.

Here, we adopt a variation of the *Asynchronous Linear Threshold* (ALT) network diffusion model [66] to capture the communication dynamics of networks that contribute to MSI. In particular, we focus on how information propagates throughout the brain, starting from different primary sensory regions (e.g. primary visual cortex, auditory cortex, and different somatosensory regions). The ALT model assumes that a “node” (brain region)^1^ becomes active when more than a fraction of the neighboring nodes it receives afferent projections from are active. We use a variation of this model with weighted connections, where the weights are based on the connection density of the projections [66]. The ALT model is simple yet it incorporates information about distances between areas (to model connection delays) and uses local information (a thresholding nonlinearity) to potentially gate the flow of information. Figure 1 illustrates the notion of *activation cascade*, showing only a small part of the visual cascade according to the ALT model. With such a model, it is possible to understand how activation cascades propagate in the brain, and then combine them to study the global architecture of MSI.

**Figure 1:**
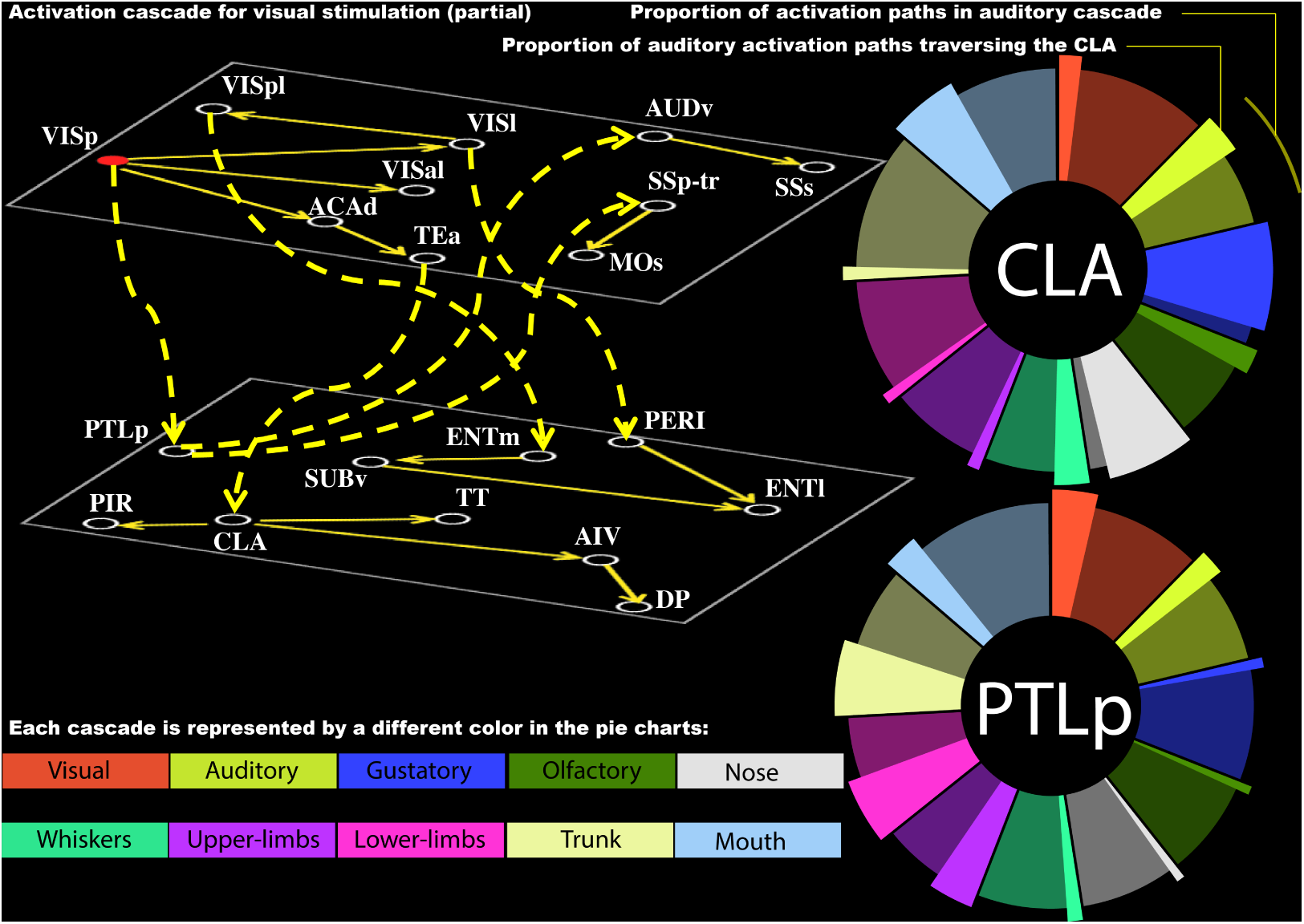
High-level illustration of approach and main result: **Top-Left:** An illustration of a single sensory cascade, originating at the primary visual cortex (VISp). The yellow edges show anatomical connections that participate in the cascade. The ROIs at the upper layer reside on the cortical surface while the ROIs at the lower layer are deeper in the brain. The edges between the two layers are dashed. To simplify the visualization, we only include 20 ROIs at this cascade (the complete cascade includes 67 ROIs). The cascade forms a Directed Acyclic Graph (DAG), and it is produced by applying the Asynchronous Linear Threshold (ALT) model on the cerebral cortex portion of the mouse connectome – starting the cascade at VISp. **Bottom and Right:** The two most important core ROIs at the hourglass waist – Claustrum (CLA) and Posterior Parietal (PTLp) cortex – jointly cover 40% of all activation paths in the ten sensory cascades we consider. Each cascade is represented by a different color. We include two circular disks (pie charts), one for each of these two core ROIs. The proportion of activation paths in each cascade is shown by the corresponding color. The protruded portion of each circular section represents those paths that traverse the corresponding core ROI. For example, the activation paths in the auditory cascase account for about 8.5% of the total number of paths – and about 36% of those paths traverse CLA.

We apply this model to the Mouse Connectivity Atlas (MCA) provided by the Allen Institute for Brain Science [42]. This connectome has been available since 2014, and it consists of estimates of connection density between cortical as well as subcortical regions, providing access to information about whole brain connectivity across functionally and structurally distinct regions. By coupling the ALT network diffusion model with this representation of the connectome, we can ask questions such as: what is the relative order in which different regions get activated after, say a visual or auditory stimulation? Which are the most central regions for each activation cascade? Are these cascades largely independent of each other, or are there few “bottleneck” regions through which almost all cascades go through? If so, which are these regions and what is their topological role in each cascade?

To examine the accuracy of the ALT model, we use Voltage Sensitive Dye (VSD) imaging to capture how the activity that is triggered from uni-sensory stimulation propagates throughout the cortex [35]. By comparing our modeling results to these functional datasets, we find that the ALT model predicts correctly the temporal ordering of node activations in a cascade – as long as the VSD data provides sufficient temporal resolution to permit activation time comparisons. This suggests, perhaps surprisingly, that despite the simplicity of linear threshold models, they can provide a useful approach for studying communication dynamics in brain networks.

We then aggregate the uni-sensory activation cascades predicted by the ALT model to investigate the architecture of MSI. To do so, we consider the number of activation paths that traverse each node across all of the different uni-sensory cascades inferred by the ALT model. We find that a small set of brain regions (around ten), form a “bottleneck” through which almost all such activation paths traverse. The *Claustrum (CLA)*, despite its small size, is the most central bottleneck region in the flow of sensory information from primary sources towards higher-level brain regions [28]. The *Posterior Parietal (PTLp)* cortex is the second bottleneck node, covering almost as many activation paths as CLA. Figure 1 visualizes the contribution of these two core regions in each of the ten sensory cascades we consider. About 40% of all activation paths, across all sensory cascades, traverse these two regions, suggesting that they play a prominent role in MSI.

Our results support the presence of a *bow-tie or hourglass architecture* in the architecture of MSI. The salient qualitative feature of an hourglass architecture is that a small number of nodes (at the waist of the hourglass) can cover almost all source-target paths [50]. In the context of MSI, this means that the multi-modal sensory input is first integrated through a small set of brain regions at the *waist* of the MSI hourglass. Then, those intermediate-level representations diverge to several higher-level association regions, providing integrated multi-sensory information for more complex cognitive tasks. Importantly, this result would not be revealed through static network analysis metrics and methods (such as betweenness centrality or rich-club), suggesting that the dynamic perspective offered by the ALT model provides valuable insights into MSI.

## 2 Methods and Data

The ALT model focuses on how stimulation of one cortical region propagates to the rest of the brain when constrained by the underlying connectome. An appropriate metaphor could be to think of the cortex as a complex consisting of several ponds that are connected through creeks of different lengths and widths (the connectome), while a stimulus corresponds to a large rock falling in one of the ponds. ALT aims to model how that initial perturbation in the pond ripples through the connected ponds (the activation cascade). In the rest of this section, we describe the main components of our approach: the mouse connectome, the ALT diffusion model, the hourglass network analysis framework, and the VSD-based model validation.

### 2.1 Structural network

The structural (i.e., anatomical) network we analyze is a subset of the Allen Mouse Brain Connectivity Atlas [42], which is based on tracking axonal projections labeled by viral tracers. It consists of 213 ROIs that cover the entire brain – all tracer injections however were performed at the right hemisphere [42]. This means that the contralateral connections from the left hemisphere to the right and the ipsilateral connections at the left hemisphere are not mapped. For this reason we only analyze the right hemisphere connections.

The strength of the connection from a source ROI to a target ROI is quantified by a metric that Oh et al. refer to as *connection density* [42]. This metric is roughly proportional to the average number of axons projecting to a target ROI neuron from the source ROI. This metric can be thought of as the total number of axons from the source ROI to the target ROI, normalized by the size of the target ROI.

Each connection is associated with a p-value that quantifies the statistical confidence that that connection exists [42]. We filter out connections with p-value higher than 0.05.^2^ The reason we do not filter connections based on their weight is because there are many weak but statistically significant connections, as shown in Figure SI-1 – it is well known that weak connections can play an important role in network diffusion phenomena as long as there are many of them [12].

The physical length of the connections is approximated based on the Euclidean distance between the two corresponding ROIs’ centroids. This is only an approximation but it is reasonably accurate because the mouse cortical surface is small and smooth with little folding [38, 60]. Additionally, as shown in Section 3.6, our results are robust to the selection of the connection weights or physical lengths.

The network we consider consists of the 67 ROIs that cover the four components of the cerebral cortex: *isocortex, olfactory areas, hippocampal formation, and cortical subplate*. We do not include subcortical ROIs that reside in the cerebellar cortex, cerebellar nuclei, striatum, medulla, pallidum, midbrain, pons, thalamus and hypothalamus for the following reasons. First, we are mostly interested in how sensory information that reaches the primary sensory regions of the cortex propagates throughout the rest of the cortex [59], rather than on the role of subcortical structures such as the thalamus or the superior colliculus. Most of the MSI literature focuses on the how those subcortical structures modulate and “route” sensory information in different parts of the cortex [30] – this type of processing however cannot be modeled without further sub-divisions of the thalamus and other subcortical regions and without at least some cell-type specificity. Second, modeling the interactions between cortical and sub-cortical regions, especially in the context of MSI, would require more elaborate communication models that can capture feedback mechanisms. Third, the accuracy of the inferred subcortical projections in the Allen mouse connectome is not expected to be as high as that of cortical projections.

The final network, denoted as *N_c_*, consists of 617 directed edges between 67 nodes – each node corresponds to one of the ROIs in our model. The density of *N_c_* is 13.9%. The distribution of edge weights is skewed, with 80% of the edges having a weight of less than 5 and few edges having a weight of up to 40. The distribution of edge lengths is almost uniform in the 1-7mm range. The diameter of the network (maximum shortest path length between any two nodes) is 7 hops, while a node is on the average about 4 hops away from any other node. The average in-degree of each node is 9.2 connections (*σ*=3.1), while the out-degree distribution has the same mean but larger variability (*σ*=8.3). Additionally, the network *N_c_* is strongly clustered, with an average clustering coefficient of 60% [10].

The ten primary sensory regions associated with the visual, auditory, gustatory, olfactory systems, as well as six somatosensory regions for different body parts (see Table SI-1), have a special role in our analysis: they are viewed as *sources* of sensory information in the cortex [4, 61]. The location of these ROIs in the Allen Mouse Brain Atlas is shown in Figure 2.

**Figure 2:**
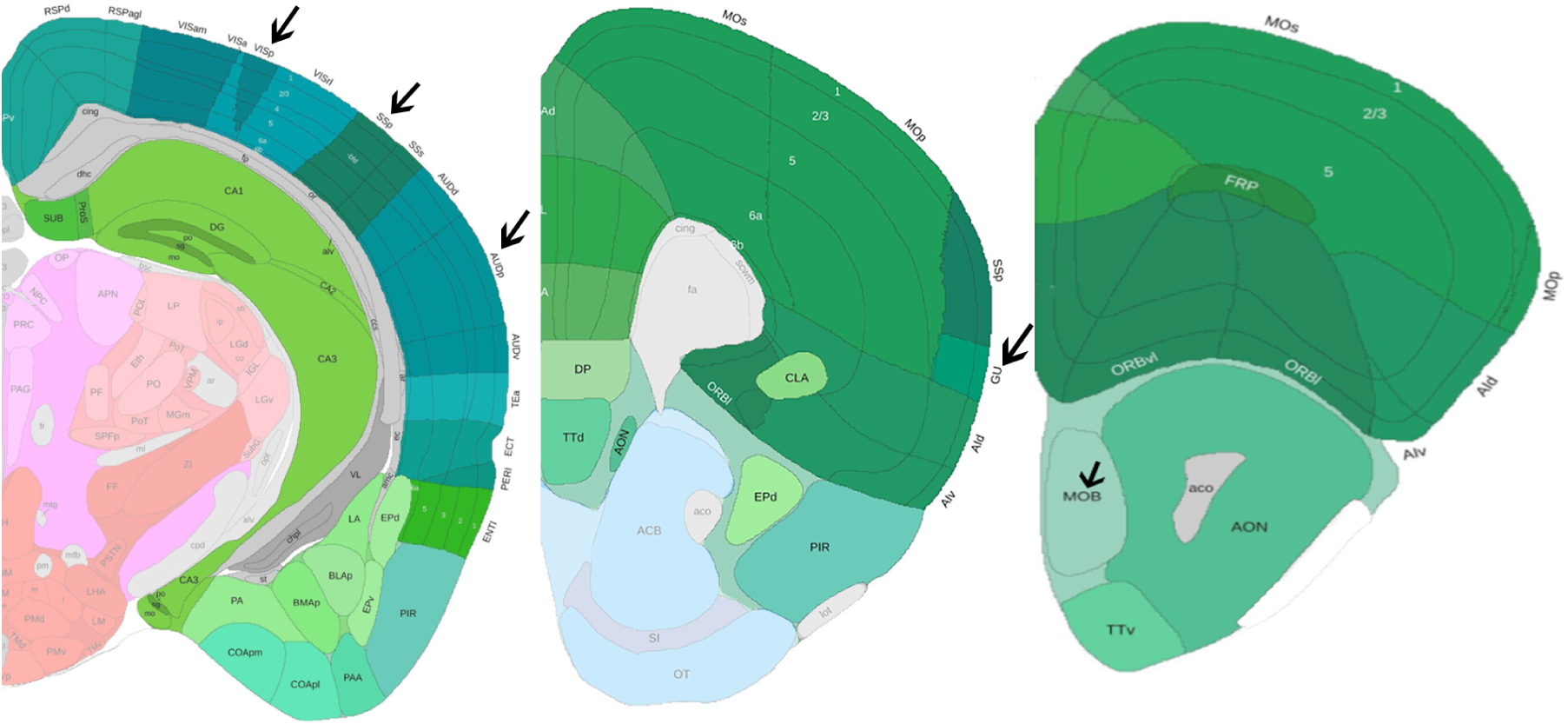
The location of the ten primary sensory regions: Three coronal slices from the Allen Mouse Brain Atlas with the cerebral cortex regions tinted by green and the source regions identified by an arrow. The somatosensory region includes six different sub-regions for lower limbs, upper limbs, trunk, mouth, nose and whiskers. The remaining four sensory sources are: visual (VISp), auditory (AUDp), gustatory (GU) and olfactory (MOB).

### 2.2 The ALT model and activation cascades

The connectome is a structural network and so it constraints, but it does not determine by itself, the paths through which information flows in the brain. To study that flow, we also need to model how dynamic brain activity propagates on the connectome.

We choose a simple network diffusion linear threshold model [15], mostly because it involves a single parameter – more realistic neural mass models, such as Wilson-Cowan, depend on several parameters [52]. The model assumes that nodes are either *inactive* (state=0) or *active* (state=1). In the model’s simplest form, a node *i* becomes active when more than a fraction *θ* of its neighbors become *active*. Here, we deploy a variation for directed and weighted networks in which each edge is associated with a communication delay and a weight, referred to as *Asynchronous Linear Thereshold (ALT)* model.

Specifically, the state of node *n_i_* is represented by *s_i_*(*t*), the neighbors of *n_i_* with an incoming edge to *n_i_* are denoted by *N_in_*(*n_i_*), the communication delay from node *n_j_* ∈ *N_in_*(*n_i_*) is *t_ji_*, while the weight of that connection is *w_ji_*. Initially, the state of every node is set to 0, except the *source* node of the activation cascade, which is set to active at time *t* = 0. The state of each node *n_i_* is then updated asynchronously based on the state of its neighbors as follows:

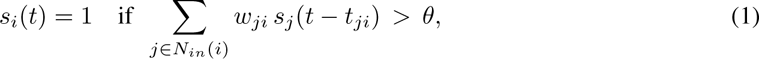

where *θ* represents the activation threshold.

Figure 3-a illustrates the ALT model with a toy example, where we can think of each node as a cortical ROI. The (directed) edges between ROIs represent the structural connections between ROIs. In this example, the activation threshold *θ* is set to 1. For this value, the cascade covers the entire network node within 7 time steps. If *θ* was larger than 1, the cascade would not take place – the only active node would be *n*_1_. As we will see in the next section, this sharp transition between not having a cascade and a complete cascade as *θ* decreases also occurs in the mouse connectome.

**Figure 3:**
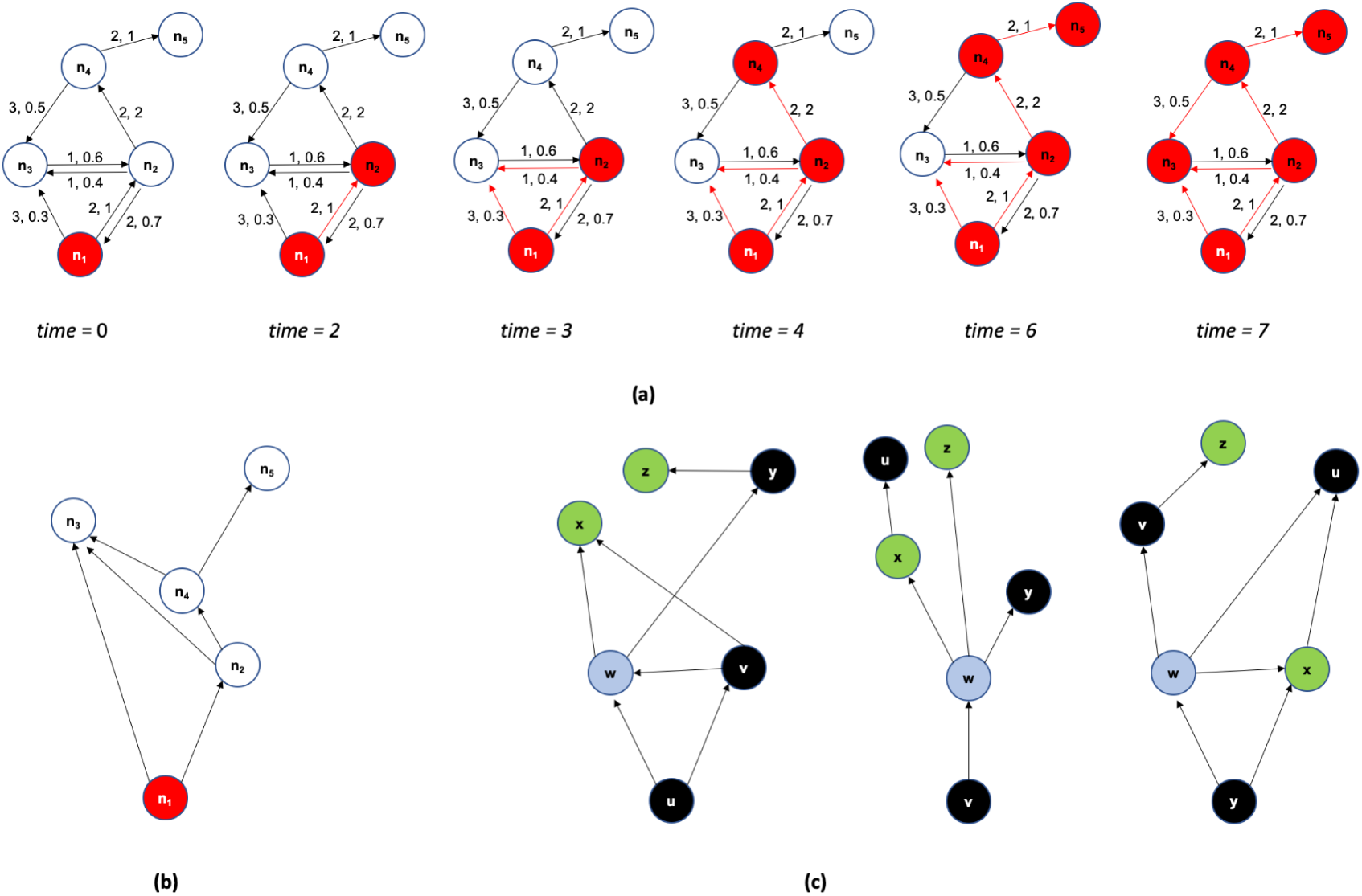
Illustration of ALT model and τ-core analysis: **(a)** A toy example of a 5-node network on which we run the ALT model. Each edge is marked with a communication delay, followed by a weight. The activation threshold is *θ*=1. The black edges represent the underlying structural network while the red unidirectional edges represent the activation cascade as it unfolds over time. **(b)** The activation cascade directed acyclic graph for the previous toy example. The source of the cascade is *n*_1_. **(c)** A toy example with three activation cascades (the sources are nodes *u*, *v* and *y*). The total number of source-target paths is 11 (4 at the left, 3 at the middle, and 4 at the right). Node *w* has the highest path centrality (*P* (*w*)=9/11). If *τ ≤* 9/11, the *τ*-core consists of only that node.

An activation cascade also reveals the node(s) that contribute in a causal manner in the activation of a node. For example, the activation of *n*_1_ and of *n*_2_ in the previous example is not sufficient to activate *n*_3_; the latter is activated only when *n*_1_, *n*_2_ and *n*_4_ are all active. Suppose that *t_i_^a^* denotes the time at which *n_i_* becomes active according to the ALT model. We say that node *n_j_* contributes in the activation of *n_i_* (denoted by *n_j_ → n_i_*) if *n_j_* ∈ *N_in_*(*n_i_*) and 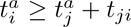. In other words, the nodes *j* that contribute to the activation of *n_i_* have a connection to *n_i_* and they should be active at least *t_ji_* time units before the activation of *n_i_*. The set of relations *n_j_ → n_i_* form a Directed Acyclic Graph (DAG) with a single source node that covers all nodes that participate in the cascade. The DAG edges are a subset of the connections in the underlying structural network. The cascade for the previous example is shown in Figure 3-b.

### 2.3 Analysis of activation cascades

An activation cascade consists of a collection of *source-target* paths, with each such path originating at the source of the cascade and terminating at a node without any outgoing edges in the cascade. A source-target path is a sequence of ROI activations that propagate in a causal manner from the source node to a target node. For instance, the cascade of Figure 3-b includes three source-target paths from *n*_1_ to *n*_4_ because the activation of the latter requires the activation of both *n*_2_ and *n*_3_, and the activation of *n*_2_ requires the activation of both *n*_1_ and *n*_3_.

After constructing a cascade for each source, we use network analysis to identify the nodes that play a more central role in the collection of all cascades. The centrality metric we use has the following graph theoretic interpretation: for each node *v*, the *Path Centrality P* (*v*) of node *v* is the fraction of all source-target paths, across all cascades, that traverse *v* (following the terminology of [50]). Figure 3-c illustrates this metric with three small cascades. Nodes with higher path centrality (PC) are expected to be more important because the activation cascades depend more heavily on them. Note that a source node is traversed by all source-target paths in its own cascade but it may have low path centrality when we consider the collection of all cascades.

The PC metric quantifies the importance of each node in isolation. We are interested, on the other hand, in the smallest set of nodes that can jointly cover almost all source-target paths in the given set of cascades. To answer this question, we adopt the *τ-core* definition of [50]: *the τ* **-core** *is the minimal set of nodes such that the fraction of source-target paths that traverse any node in the set is at least τ.* The *τ*-core problem is NP-Hard for *τ <* 1 [50]. It can be approximated with a greedy heuristic in which the node with the highest PC joins the set in each iteration. That node is then removed from all cascades it appears in. The PC of the remaining nodes is recomputed after each iteration. If the *τ*-core is a small set, relative to the total number of nodes in the network, the nodes of that set can be thought of as the *bottleneck* of the activation cascades (see Figure 3-c for an example).

### 2.4 Comparison of modeling results with functional data

To examine whether the ALT model can accurately predict the propagation of sensory stimulation in the cortex, we need an experimental setup in which we can stimulate different sensory modalities of a living animal, and monitor at the same time and in a fine temporal resolution (in the order of a millisecond) the activity of different cortical neural populations (in a spatial resolution of few *µ*m).

Such experiments are possible today, relying on technologies such as calcium imaging or voltage-sensitive dies (VSD) in conjuction with fluorescence microscopy. Here, we leverage the experimental results of an earlier study to examine the accuracy of the ALT model in the context of whole-cortex imaging in mice under single sensory stimulation [35]. In brief, the experiments include five types of sensory stimulation: visual (flash), auditory (tone), whisker touch, forelimb touch, and hindlimb touch. Each stimulation experiment is repeated ten times and on several different animals (we analyze data for five animals). The recorded images cover almost the entire cortical surface, have a temporal resolution of 6.67msec, and a size of 128 *×* 128 pixels at a spatial resolution of 50*µ*m/pixel. As it will become clear in Section 3.2, the experimental spatial resolution is sufficient for our purposes, given that each network node in the ALT model refers to an entire cortical ROI of the Allen mouse brain atlas. The temporal resolution, however, is marginally sufficient because for about 20-30% of all ROI pairs we cannot tell which ROI gets activated first as they appear to be activated during the same frame.

To compare the VSD-based results with our modeling results, we perform the following steps:

**Step 1. Register:** Register the native cortical surface of each animal to the Allen mouse brain atlas. This step is performed using an affine transformation that minimizes the least-squares error between the coordinates of the centroid of a primary sensory ROI (e.g., VISp) in the Allen atlas, and the coordinates of the pixel that first gets activated after the corresponding sensory stimulation (visual in this example). We use five primary sensory ROIs to construct this transformation: VISp, SSp-bfd, SSp-ll, SSp-ul and AUD-p.

**Step 2. Parcellate into ROIs:** Parcellate the native cortical surface into ROIs using the Allen mouse brain atlas and the previous affine transformation. Some cortical ROIs are not visible in the VSD images: FRP, PL, ACAd, VISpl, and VISpor. Also, the MOs region is only partially captured in the VSD images.

**Step 3. Estimate Activation Time:** Estimate an *activation time* for each ROI in the experimental data. To perform this step, we first identify the maximum post-stimulus activity for each pixel in that ROI – this is defined as the activation time of that pixel. The image frame that corresponds to the activation time for most pixels of that ROI is defined as the activation time of the ROI. This processing pipeline is summarized in Figure 4.

**Figure 4:**
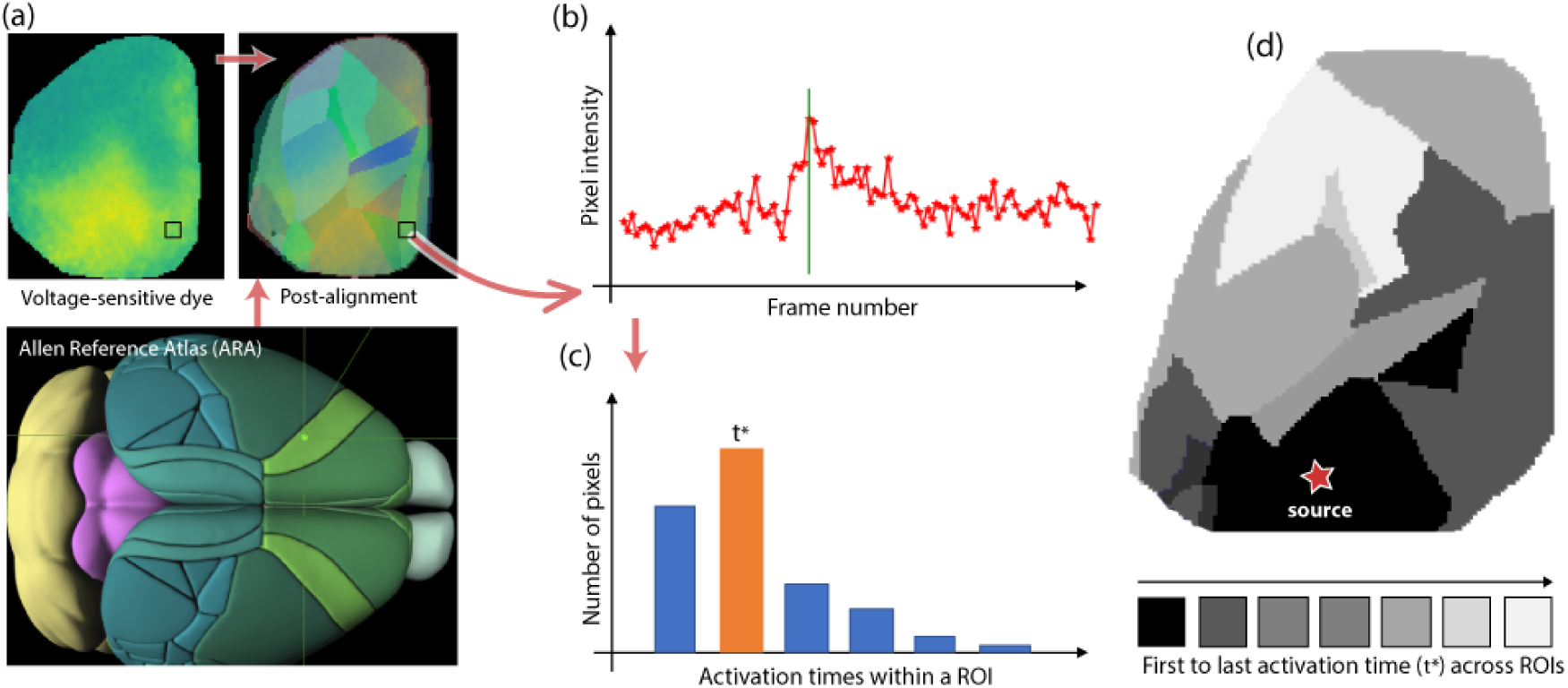
VSD data processing pipeline: (a) *Lower:* The Allen Reference Atlas (ARA). *Upper-left:* A sample VSD image covering most of the left cortical surface five frames after visual stimulation. *Upper-right:* The ROIs at the left ARA cortical surface mapped to the native cortical surface of an animal. (b) The activation time of a pixel is defined as the frame of maximum post-stimulus VSD signal at that pixel. (c) The activation time of an ROI is defined as the activation time of *most* pixels in that ROI. (d) The output of this pipeline is an activation time for each ROI, depicted here with a grey-scale map (black for the first ROI activation and white for the last).

The ALT model, on the other hand, models each cortical ROI as a single node, and it assumes that the transition of each node from inactive to active occurs instantaneously. The time unit in the ALT timeline is arbitrary – so we cannot just compare the absolute activation times between the experimental and modeling results. Instead, we examine the consistency of the temporal ordering of activations in the ALT model and in the VSD experiments. Specifically, if *X* and *Y* are two ROIs, and X is activated before Y in the modeling results, is it also the case that *X* is activated before *Y* in the VSD data? If so, we count that ROI pair as a *temporal agreement*. If *X* and *Y* are activated in the same VSD frame, we count that pair as a case of *insufficient temporal resolution*. Finally, if *X* is activated after *Y* in the VSD data, we count that ROI pair as *temporal disagreement*.

## 3 Results

### 3.1 Sensory-specific activation cascades in the mouse cortex

The ALT model requires the selection of a single parameter, the activation threshold *θ*. This threshold controls the *size of the cascade*, i.e., the fraction of network nodes that become active after the activation of a source node. One may expect that as *θ* decreases towards zero, the size of the cascade becomes *gradually* larger. This is not the case however. Figure 5-a shows that as *θ* decreases we observe a rapid transition from the absence of a cascade (where only the source node is active) to a complete cascade, in which all nodes become active. This is true for all source nodes listed in Table SI-1.

**Figure 5:**
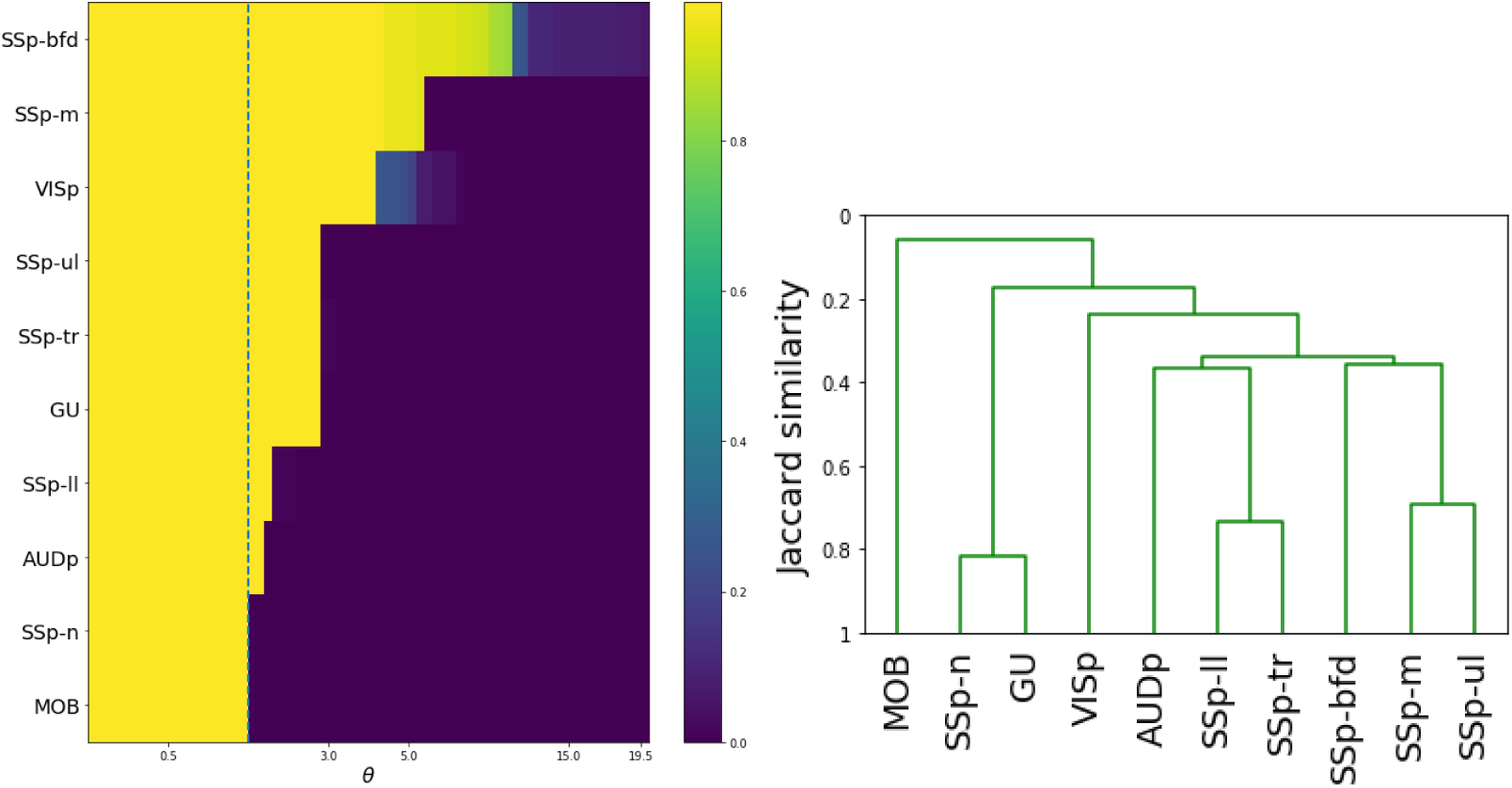
Effect of parameter θ on cascade size, and similarity between the ten cascades: (a) Each row of the heat map shows the fraction of activated nodes after the stimulation of a single source, for different values of the threshold *θ*. The selected threshold is marked with the dashed vertical line. (b) Similarity between the ten sensory cascades using the average-linkage hierarchical clustering method.

The reason behind this rapid transition is the highly clustered topology of the mouse connectome. This property is often quantified by the *clustering coefficient* [39], which has an average value of 0.60 in the mouse connectome. This means that if a node *v* connects to two nodes *u* and *w*, there is a probability of 60% that there is an edge from *u* to *w* or in the opposite direction,“closing the triangle” formed by the three nodes *u, v, w*. So, if *v* is the source of the activation and *θ* is low enough so that *v* activates at least one of its neighbors, say *u*, it is highly likely that other neighbors of *v* receive input from *u* as well, increasing the chances that they will also get activated. The same argument applies to all other pairs of activated nodes – not only the source and its direct neighbors.

We choose *θ* so that the ALT model produces a *complete cascade*, for every source we consider [3]. This choice is motivated by experimental results [35]: at least some activity is detected in all cortical regions after sensory stimulation (visual, auditory, touch) in anesthetized mice. It may appear counter-intuitive that visual stimulation, for example, can impact activity in regions associated with different sensory modalities (e.g., gustatory) but such interactions are possible through the many feedback connections in the connectome, and they are consistent with several prior studies which argue that there are no strictly uni-sensory regions, and that to some extent the entire cortex is a multi-sensory organ [5, 11, 13, 37].

Lower values of *θ* would also result in complete cascades. However as *θ* decreases, the dynamics of the underlying cortical networks would move away from the critical boundary between sensitivity to internal or external stimulation and stability [45]. A very low value of *θ* would keep the system in a constant state of activation, resembling a brain under a global epileptic seizure.

Based on the previous considerations, we select *θ* = 0.98 as *the lowest value that results in a complete cascade across all sensory modalities.*^3^ We have repeated the analysis for two more values of *θ* (0.90 and 0.95) without any significant changes in the results (see Section SI.4).

Figure 6 shows the complete activation cascade when the source of the stimulation is the primary visual cortex (VISp) – the corresponding cascades for the other sensory modalities are included in the Supplementary Information (see Section SI.3). Note that the activation of the source triggers the activation of eleven other ROIs. Only few of them however play a major role in extending the cascade to the rest of the network: ECT (ectorhinal), PTLp (posterior parietal association), VISl (lateral visual), and POST (postsubiculum). PTLp in particular, causes the activation of seven more ROIs at the next step of the cascade. The activation of the claustrum (CLA), in this cascade, takes place through the sequence of ECT, followed by TEa (temporal association).

**Figure 6:**
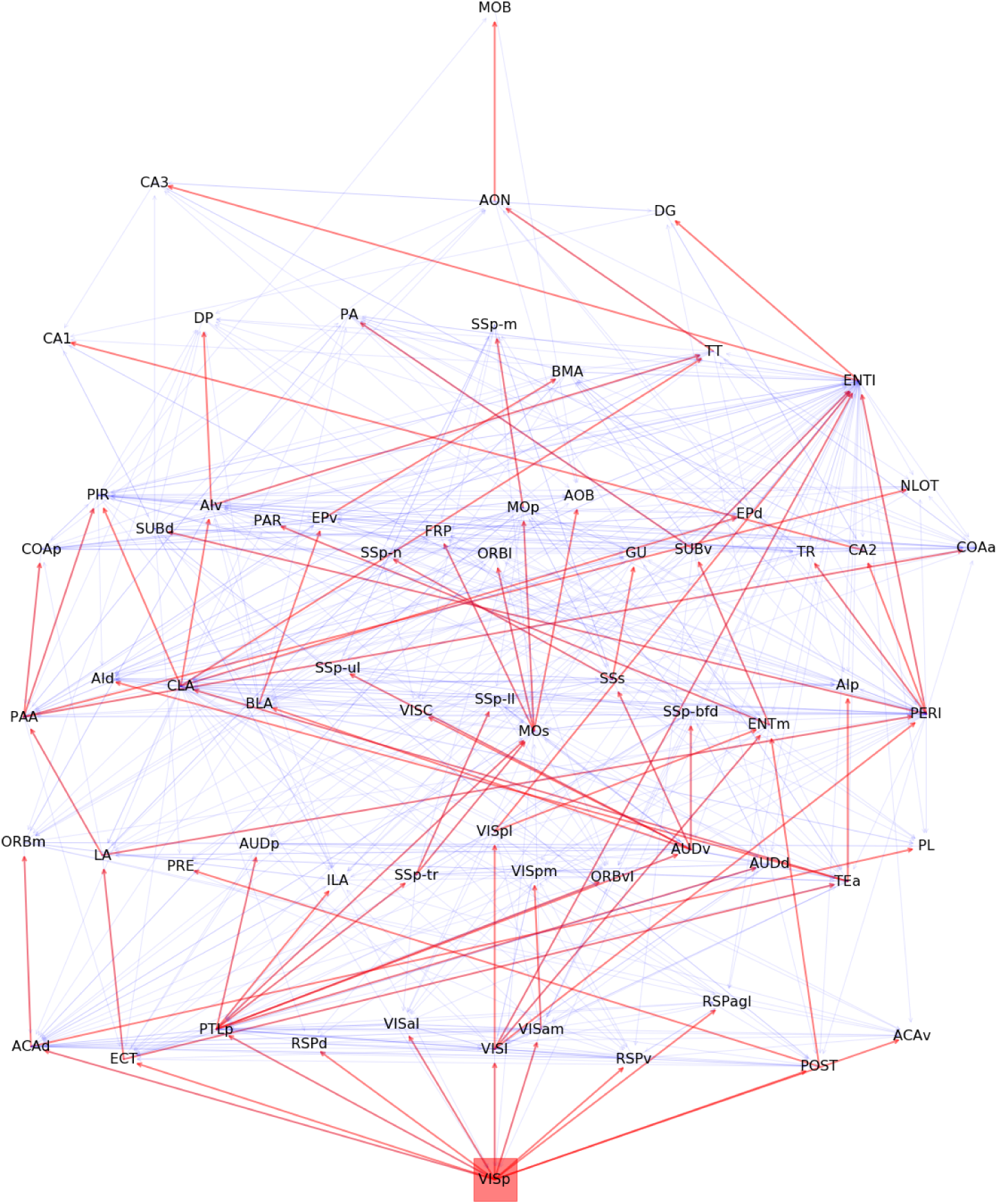
The visual activation cascade, accoring to ALT: The source for this cascade is the primary visual cortex (VISp). The red edges form the activation cascade, while the underlying grey edges show anatomical connections that do not participate in this cascade – those connections may be present in other sensory cascades or they may play a role in feedback (or second-order) interactions that are not captured by the “first ripple” scope of the ALT model. To help with the visualization, we place the nodes in eight layers, so that cascade edges only point from a layer to a higher layer (never to the same or lower layer). The vertical position of each node is slightly “jittered” to avoid cluttering due to anatomical connections between nodes of the same layer.

We emphasize that the hierarchical layout shown in Figure 6 is specific to each sensory modality and it represents the activation cascade from the corresponding source to the rest of the cortex. This notion of hierarchy should not be confused with the hierarchical organization of the cortex [26] that results from anatomical distinctions of intracortical connections (feedforward versus feedback, based on laminar markers) [19]. The latter is an anatomical hierarchical structure, it is not specific to any particular sensory modality, and it does not convey any functional information about how one ROI may be affecting another in the presence of a specific external or internal stimulation. An activation cascade, on the other hand reveals the sequence and causal dependencies through which ROIs get activated after an initial activation at the source ROI.

### 3.2 Model validation

We examine the validity of the ALT model predictions using functional imaging data during sensory stimulation experiments, as discussed in Section 2.4. The question we focus on is: *after stimulating a specific sensory modality (e.g., visual), if the ALT model predicts that an ROI X should be activated before an ROI Y, is it true that X gets activated before Y in the functional imaging data?* When this is the case, we count the pair (*X, Y*) as a *temporal agreement*. If *X* gets activated before *Y* in the ALT model but the opposite is true for the experimental data, we count (*X, Y*) as a *temporal disagreement*. Because of the finite temporal resolution in the experimental results (each frame is sampled every 7msec roughly), there are also cases where *X* and *Y* appear to be activated during the same frame, while the model always predicts a temporal difference between two activations – when that is the case, we count (*X, Y*) as a case of *insufficient temporal resolution*.

Figure 7a shows the percentage of (*X, Y*) ROI pairs that show temporal agreement, temporal disagreement, and insufficient temporal resolution between the activation order of *X* and *Y* in the modeling results and the mouse experiments. The plot shows results for five animals and for five sensory stimulations. Even though the variability across animals is considerable, we observe that the percentage of temporal agreement pairs, averaged across the five animals, is higher than 50% and it varies between 55% to 80% depending on the sensory modality. On the other hand, the corresponding percentage of temporal disagreement is less than 10%-20%, depending on the stimulation. In the rest of the cases, the temporal resolution is not sufficient.

**Figure 7:**
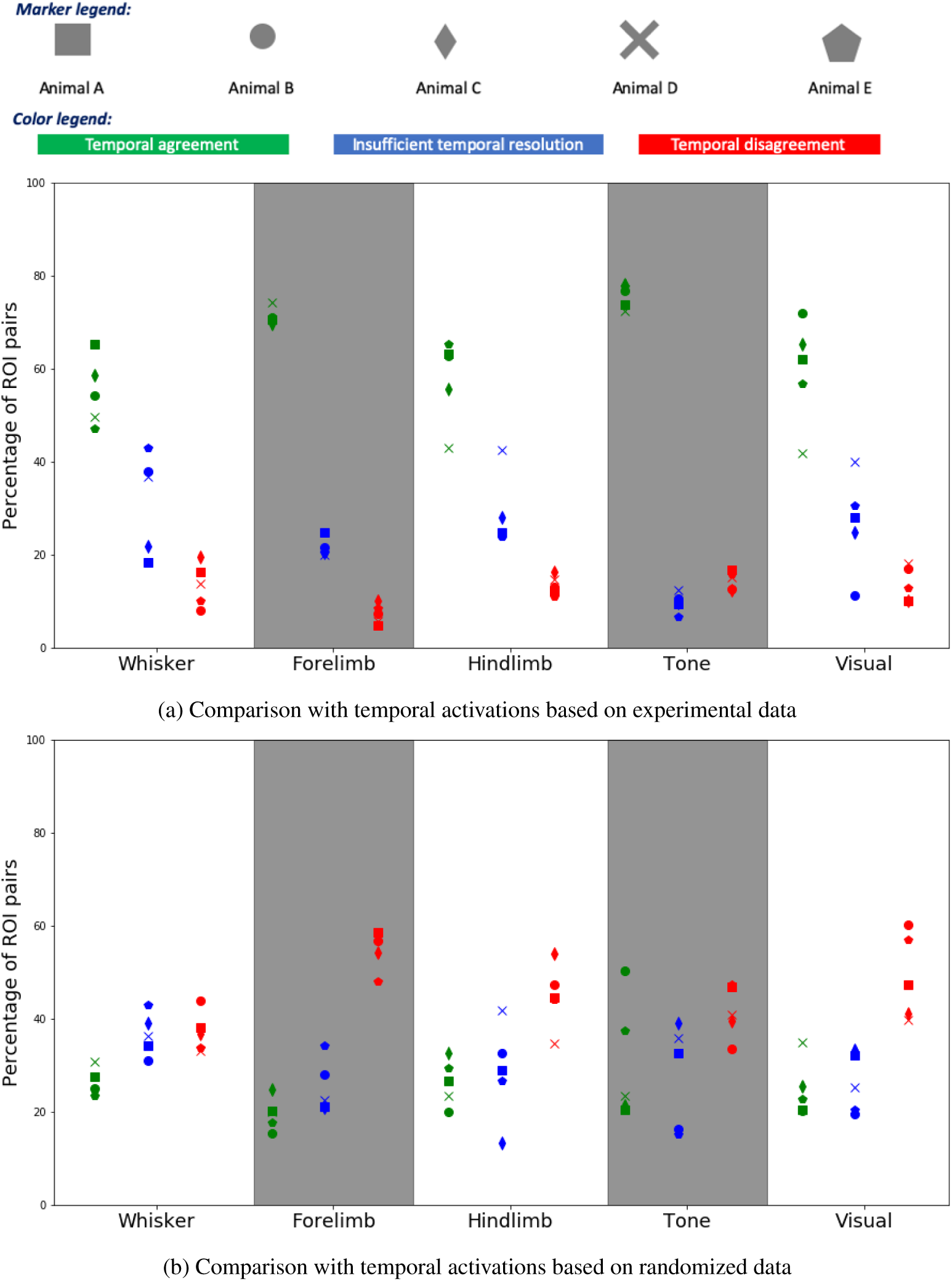
Comparison between model-based and experimental temporal ordering of ROI activations: a) The y-axis shows the percentage of (*X, Y*) ROI pairs that show temporal agreement (green), temporal disagreement (red), and insufficient temporal resolution (blue) between the activation order of *X* and *Y* in the modeling results and the mouse experiments. The plot shows results for five animals and for five sensory stimulations (a touch at the whiskers, forelimb, and hindlimb, as well as an auditory and a visual stimulation). b) The same comparison but here we have randomized the ROIs that are active during each frame, preserving the number of ROI activations in each frame.

We also compare the ALT modeling results with a randomized sequence of experimental activations in which we preserve the number of ROIs that are activated in each frame but assign random ROI activations during each frame. The results of that experiment are shown in Figure 7b. The average percentage of ROI pairs that show temporal agreement in this case varies between 20%-30%, depending on the stimulation. This comparison shows that the ALT model has significant prediction power on the temporal sequence of ROI activations, relative to a randomized baseline.

Finally, we analyzed the temporal disagreement cases between VSD experiments and modeling results to examine if certain brain regions, or pairs of regions, are over-represented in those disagreements (see Section SI.5). The main result of that analysis is that the ROIs with highest disagreement cases appear at the boundary of the cortical surface at the VSD datasets and they are only partially visible. So, it is likely that the VSD data may not accurately capture the time at which those boundary regions are activated after each stimulation.

### 3.3 Similarity of sensory cascades

We next sought to answer the question of how similar the ten sensory cascades predicted by the ALT model are. The similarity between two cascades can be quantified using the *Jaccard similarity* metric. It is defined as the ratio of the common connections in two cascades over the total number of connections in those cascades.

After calculating the Jaccard similarity between every pair of cascades, we use an agglomerative hierarchical clustering algorithm to construct a dendrogram of the ten cortical sensory cascades. This dendrogram was computed for three linkage methods: *average linkage* (the similarity between two clusters is based on the average similarity across all pairs of cascades in the two clusters), *single linkage* (based on maximum similarity), and *complete linkage* (based on minimum similarity). The resulting dendrograms are quite similar across the three linkage methods. Figure 5-b shows the dendrogram with average linking – the two others are shown in Figure SI-13.

A first observation is that the olfactory cascade (originating at MOB) is very different than all other sensory cascades – its similarity is less than 10% with the cluster of all other cascades. This is expected given that olfaction is quite different than all other sensory processes – it bypasses the thalamus and MOB is the only primary sensory ROI in the mouse connectome that is not located in the isocortex [23, 56].

Interestingly, the two most similar cascades are the Gustatory (GU) and the somatosensory cascade of the nose (SSp-n). Further, these two cascades are quite different than all others, including the rest of the somatosensory cascades. The somatosensory cascades are quite similar to each other and they tend to cluster as follows: trunk and lower-limb (similarity of about 75%), mouth and upper-limb (about 70%), while the whiskers produce a significantly different cascade than the previous four (similarity of about 40%). This organization mirrors the anatomical layout of the somatosensory regions. The auditory and visual cascades are also quite distinct from all other cascades – but not as dramatically different as olfaction.

### 3.4 Core ROIs and hourglass architecture

In this section, we analyze the collection of ten activation cascades (one cascade for each source) using the network analysis approach described in Section 2.3. The total number of source-target paths in the ten cascades is 560. The path centrality distribution, which captures how many activation paths traverse each node, is shown in Figure 8a. Almost half of the nodes have very low path centrality (2% or less). On the other hand, there are four nodes with much larger path centrality – each of them covering about 20% of the source-target paths in the collection of activation cascades. These ROIs are the CLA (claustrum), SSs (supplemental somatosensory), PTLp (posterior parietal association), and AUDv (ventral auditory) areas.

**Figure 8:**
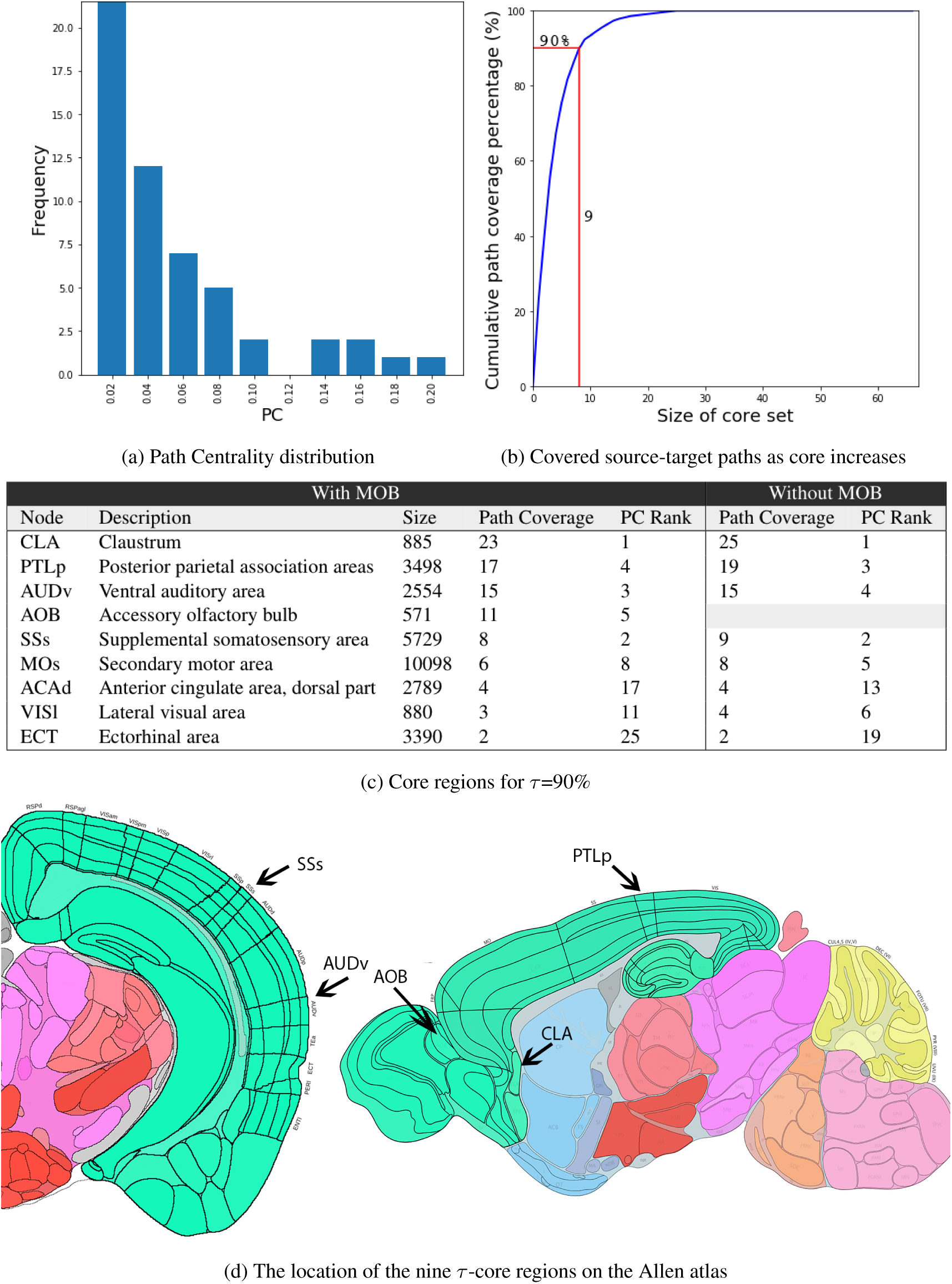
Path centrality and τ-core analysis: (a) Path Centrality (PC) histogram for the 67 regions in *N_c_*, considering all source-target paths across the ten activation cascades. (b) Cumulative path coverage by the top-X core nodes for X=1···67. Nine regions are sufficient to cover *τ* = 90% of all paths.

Some activation paths can traverse more than one of these highly central regions. For this reason, we compute *the minimal set of nodes that cover a given fraction τ of all source-target paths*, i.e., what is referred to as *τ-core* [50]. Figure 8b shows the fraction of covered source-target paths as we increase the size of the *τ*-core. The “knee-shaped” shape of this curve suggests that a small set of nodes is sufficient to cover almost all source-target paths in the activation cascades, forming a bottleneck in the MSI process. For instance, a set of nine nodes is sufficient to cover more than 90% of all source-target paths. These nine ROIs account for 14.7% of the brain volume of the ROIs we consider in the network *N_c_*.

The small size of the *τ*-core, relative to the size of the network, suggests that the cortex follows an *hourglass architecture*, in which the sensory information from all different modalities is first *integrated* (other terms could be “encoded” or “compressed”) by a small set of *τ*-core ROIs that form the “waist” (or bottleneck) of the hourglass. Those *τ*-core ROIs then drive a large number of downstream ROIs that presumably operate on multi-sensory information and contribute in higher-level cognitive processes. The benefit of an hourglass architecture is that it reduces the dimensionality of the input, computing a more compact intermediate-level sensory representation at the “waist” of the hourglass. That intermediate representation is then be re-used in more than one higher-level ROIs and cognitive tasks. The hourglass architecture is visualized, at an abstract level, in Figure 9.

**Figure 9:**
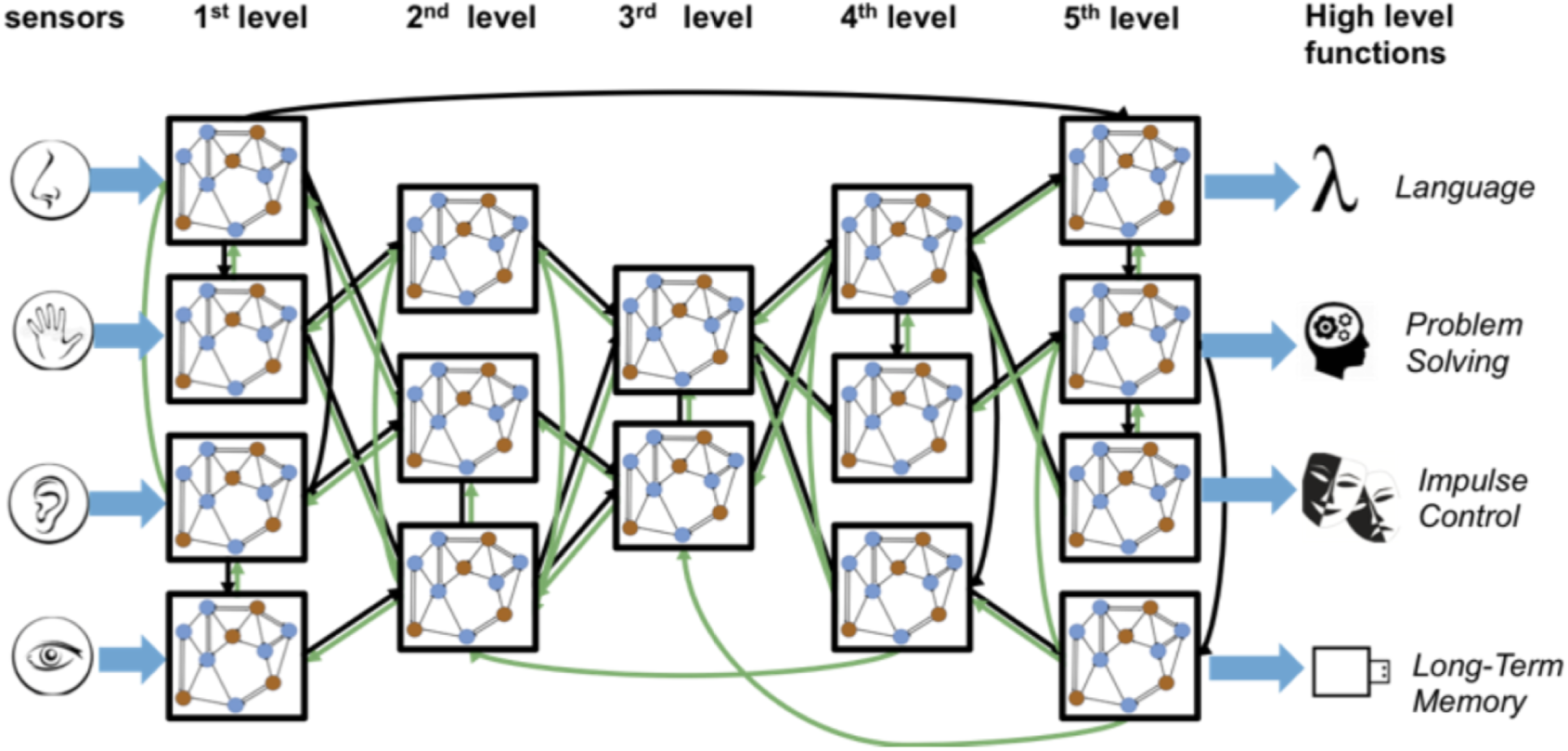
Illustration of hourglass architecture: A hypothetical network with feedforward, feedback and lateral connections between regions at different levels of the cortical hierarchy. Input information is provided at sensory-specific modules (left), while high-level cognitive tasks are performed by association regions at the other end of the hierarchy (right). The “hourglass feature” refers to the fact that the high-dimensional input information is first integrated through through a relatively small number of highly central intermediate-level regions, before it is re-used at high-level cortical regions and tasks.

The *τ*-core nodes for *τ* =90% are listed in Table 8d. Together with the percentage of *additional* source-target paths that each node contributes to the *τ*-core (“Path Coverage”), the table also shows the Path Centrality (PC) rank of that node. As expected, the node with the highest PC is the first node in the *τ*-core. After that point, the order in which nodes join the *τ*-core does not follow their PC ranking. The top three nodes (CLA, PTLp, AUDv) are sufficient to collectively cover about 60% of the activation paths. Note that none of these *τ*-core ROIs are a primary sensory region (i.e., a source node). Instead, they are either ROIs that are not typically associated with a single sensory modality (CLA, PTLp, ACAd, ECT) or they are ROIs that are often thought of as “secondary” or “supplemental” to a certain sensory modality (AOB, AUDv, SSs, MOs, or VISl). If we exclude the MOB cascade, the only difference is that the *τ*-core will not include the AOB region.

### 3.5 Location of ***τ***-core nodes in activation cascades

#### 3.5.1 Location relative to sources

In this section, we first investigate the *topological location* (rather than anatomical location) of the *τ*-core nodes relative to the source of each activation cascade. Are the *τ*-core nodes, which form the waist of the hourglass architecture, closer to the sources or targets of the activation cascades? And how does their location compare to the location of the sources and of other cortical ROIs? These questions are related to recent experimental work suggesting that cross-modal representations are constructed at early stages of the sensory information flow [6].

We focus on the top-4 *τ*-core nodes (CLA, PTLp, AUDv, SSs), which collectively cover about *τ* =70% of all source-target paths (see Table 8b). Figure 10a visualizes in grey-scale the location of each node (matrix row) in each activation cascade (matrix column). Each source node (represented with red) is obviously at a distance of zero in its own activation cascade. Note however that source nodes can be at a much larger distance from sources of other activation cascades. For instance, the primary visual cortex (VISp) appears at the maximum distance in the nose (SSp-n) and gustatory (GU) cascades.

**Figure 10:**
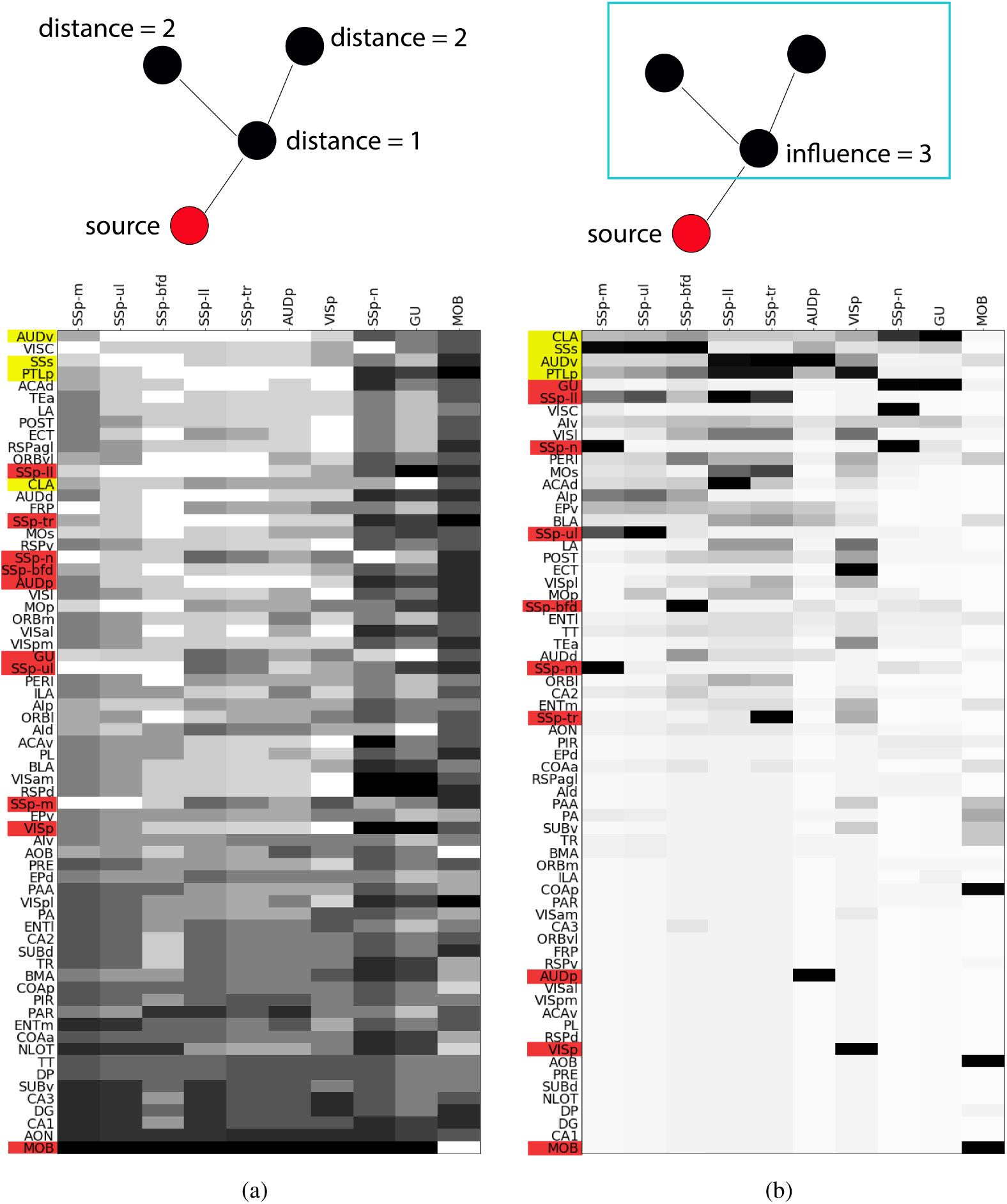
Location-related metrics: In both matrices, a column represents one of the ten activation cascades, originating at the node shown at the top of the column. (a) Each row represents the *source-distance* of the corresponding node from the source of that column’s cascade. White denotes a distance of one hop, while black denotes the maximum distance for that cascade. Rows are ordered in terms of the average distance (in number of hops) of the corresponding node from the sources of all activation cascades (excluding the MOB cascade, which is very different). (b) Each row represents the *influence* of the corresponding node, i.e., the number of nodes that are reachable from that node in the activation cascade that the column represents. White denotes an influence of one (only that node), while black denotes an influence that covers all network nodes. Rows are ordered in terms of the average influence of the corresponding node across all activation cascades (excluding the MOB cascade).

The four *τ*-core nodes we consider (represented with blue rows) *are relatively close to all source nodes*: AUDv has the lowest average distance to all of the primary source areas, while SSs and PTLp are ranked as third and fourth. The claustrum (CLA) is slightly further away from the sources, ranked 13th (out of 67) in the previous ranking. If we consider a higher value of *τ* =90%, the additional *τ*-core nodes (MOs, ACAd, VISl, and ECT) are ranked as 17th, 5th, 22nd, and 9th in terms of their average distance from sources. In summary, *all τ-core nodes appear in the top one third of the distance ranking, and so they are closer to the sources of the hourglass architecture than to its targets*.

#### 3.5.2 Location relative to targets – influence

Another way to examine the location of a node *v* in the hourglass architecture is in terms of how many nodes appear in activation paths downstream of *v* – a metric that we refer to as the *influence* of *v*. Figure 10b visualizes in grey-scale the influence of each node (matrix row) in each activation cascade (matrix column). The source of a specific cascade has, by definition, maximal influence (i.e., all network nodes) in its own cascade – but it may have a much lower influence in other cascades. Indeed, the influence of source ROIs (shown in red) does not seem to follow a coherent pattern: the gustatory (GU) and somatosensory area of the lower-limb (SSp-ll) are sources with high influence but the primary visual cortex (VISp), the primary auditory cortex (AUDp) or the main olfactory bulb (MOB) are sources with low influence in other cascades.

On the other hand, the four most important *τ*-core nodes (CLA, PTLp, AUDv and SSs) also occupy the top-four positions in terms of influence. The next four *τ*-core nodes (MOs, ACAd, VISl, and ECT) have high influence as well, ranked as 12th, 13th, 9th and 20th, respectively.

Combining the previous observations about the influence of *τ*-core nodes as well as their distance from sources, we can summarize our findings as follows: *τ-core nodes are close to most sensory sources and they also influence the activation of many downstream nodes.* These two features place *τ*-core nodes at a location that allows them to both integrate sensory information from different sources as well as to use that integrated information in driving many downstream ROIs.

### 3.6 Robustness of ***τ***-core nodes

In this section, we examine the robustness of the previous results regarding the *τ*-core when we randomize the edges and weights of the underlying connectome. We also examine whether the length and/or weights of these connections are responsible for the hourglass effect and for the specific regions that form the *τ*-core.

We create ensembles of random connectomes, derived from the mouse connectome in four different ways:

1. Randomize the weight assigned to each each edge, reallocating the weights of the original connectome across randomly selected connections but maintaining the topology.
2. Randomize the physical length of each edge (and thus its communication delay in the ALT model), again reallocating randomly the lengths of the original connections but maintaining the topology.
3. Randomize both the weights and lengths assigned to each edge, as previously mentioned. We do not maintain any correlation between weights and lengths.
4. Randomize the connectome’s topology by swapping connections between randomly selected pairs of nodes. This randomization method preserves the in-degree and out-degree of each node.

Figures 11a-11d focus on the first three randomization methods: weights, lengths, and their combination. In all cases, the *τ*-core size of the original network is contained in the 5% confidence interval of 100 randomized networks. In other words, *the weights and physical lengths of the connectome’s connections do not play a significant role in the number of τ-core nodes, for any value of τ*.

**Figure 11:**
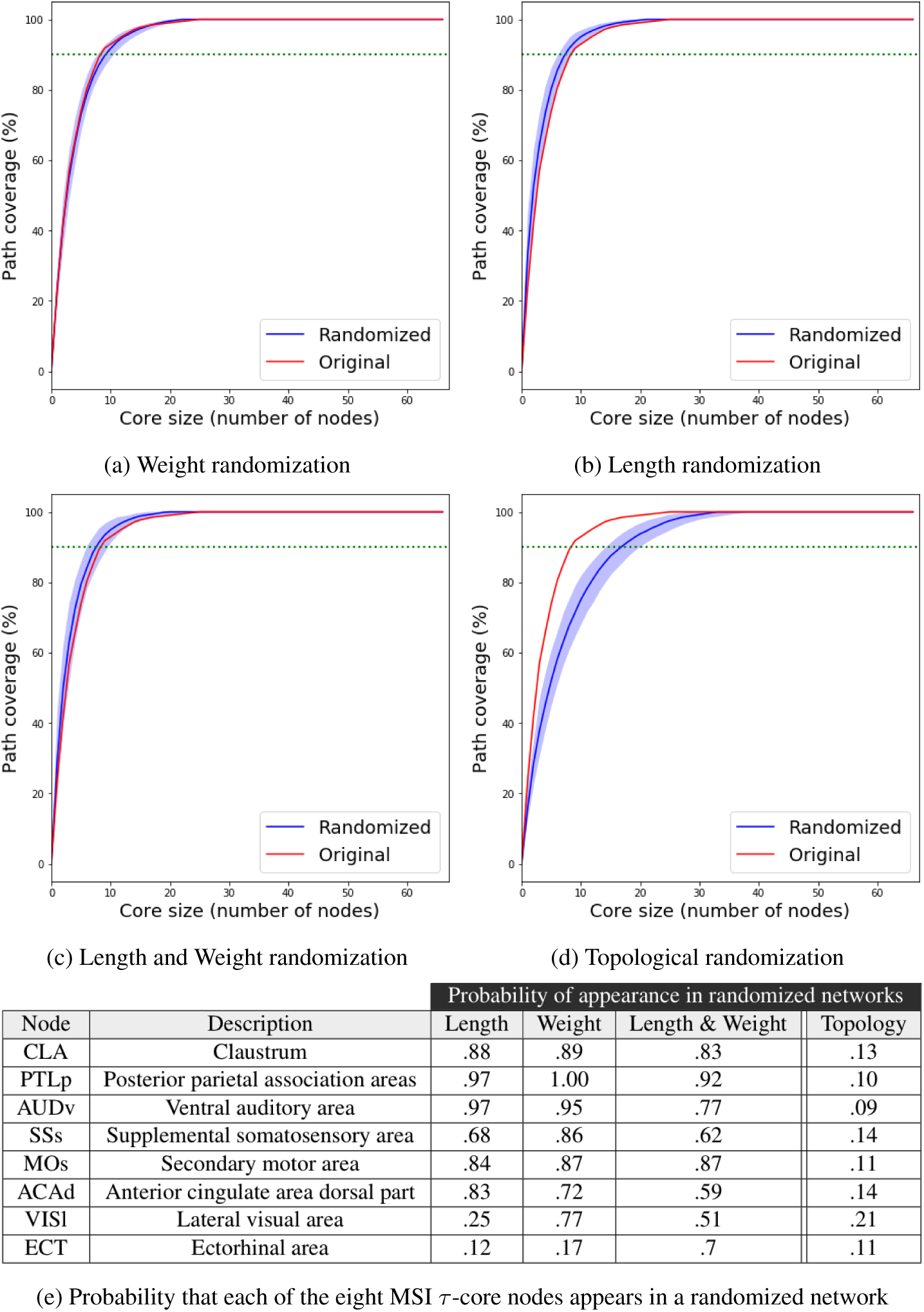
Robustness results: The effect of different connectome randomization methods on the core size. Light blue shade marks the 5% to 95% values among 100 randomization runs, while the solid blue line is the median of these runs. The red line represents the *τ*-core size for the original connectome. The dotted green line marks the *τ*-core size for *τ* =90%. The table at the bottom shows the fraction of random networks that include each of the eight *τ*-core nodes.

On the other hand, when we randomize the topology of the connectome, the *τ*-core size doubles in size when *τ* =90%: from nine nodes in the original network to eighteen. Additionally, it takes about half of the entire network to cover all activation paths in the collection of ten A-DAGs. So, *it is the graph structure of the connectome (i.e., its topology) that leads to a small τ-core size – not the weight and/or length of the connections*.

Even though the weight and length of the connections do not have a strong effect on the *τ*-core size, do they affect the identity of the ROIs that participate in that *τ*-core? To answer this question, Table 11e shows the fraction of random networks that include each of the eight *τ*-core nodes in Table 8d. The Claustrum (CLA), for instance, appears in the MSI *τ*-core of 88% of the networks that have randomized connection lengths (89% for randomized weights) but in only 13% of the networks that have randomized topology. The results are similar for the top-6 MSI *τ*-core nodes: they appear in the MSI *τ*-core of most randomized networks when we randomize connections weights and/or lengths – but they rarely appear in the MSI *τ*-core when we randomize the topology. For the last two MSI *τ*-core nodes (VISl and ECT) their membership in the MSI *τ*-core is *not* as robust: randomizing connection lengths has a major effect in the appearance of VISl in the *τ*-core, and randomizing any aspect of the network has a major effect in the appearance of ECT in the *τ*-core.

### 3.7 Which anatomical connections are more important in sensory cascades?

*Which anatomical connections are more important in terms of MSI?* Only about half the connections of the anatomical connectome appear in sensory activation cascades, and a quarter of the former appear in only one sensory cascade.

To answer this question, we examine the conditional probability that an connection *e_l_* of physical length *l* appears in an activation cascade given that *l > l*_0_; we denote this probability as *P*[*e_l_* = 1*|l > l*_0_]. Similarly, we define the probability *P*[*e_w_* = 1*|w > w*_0_] for an edge of weight *w*.

Figure 12 shows these two conditional probabilities separately for each of the ten activation cascades.^4^ Even though there are significant variations across the ten cascades, all of them show that *P*[*e_l_* = 1*|l > l*_0_] decreases with *l*_0_, i.e., *as the length of a connection increases (and especially when l is larger than 1-2mm) it becomes less likely that it will be part of an activation cascade*.

**Figure 12:**
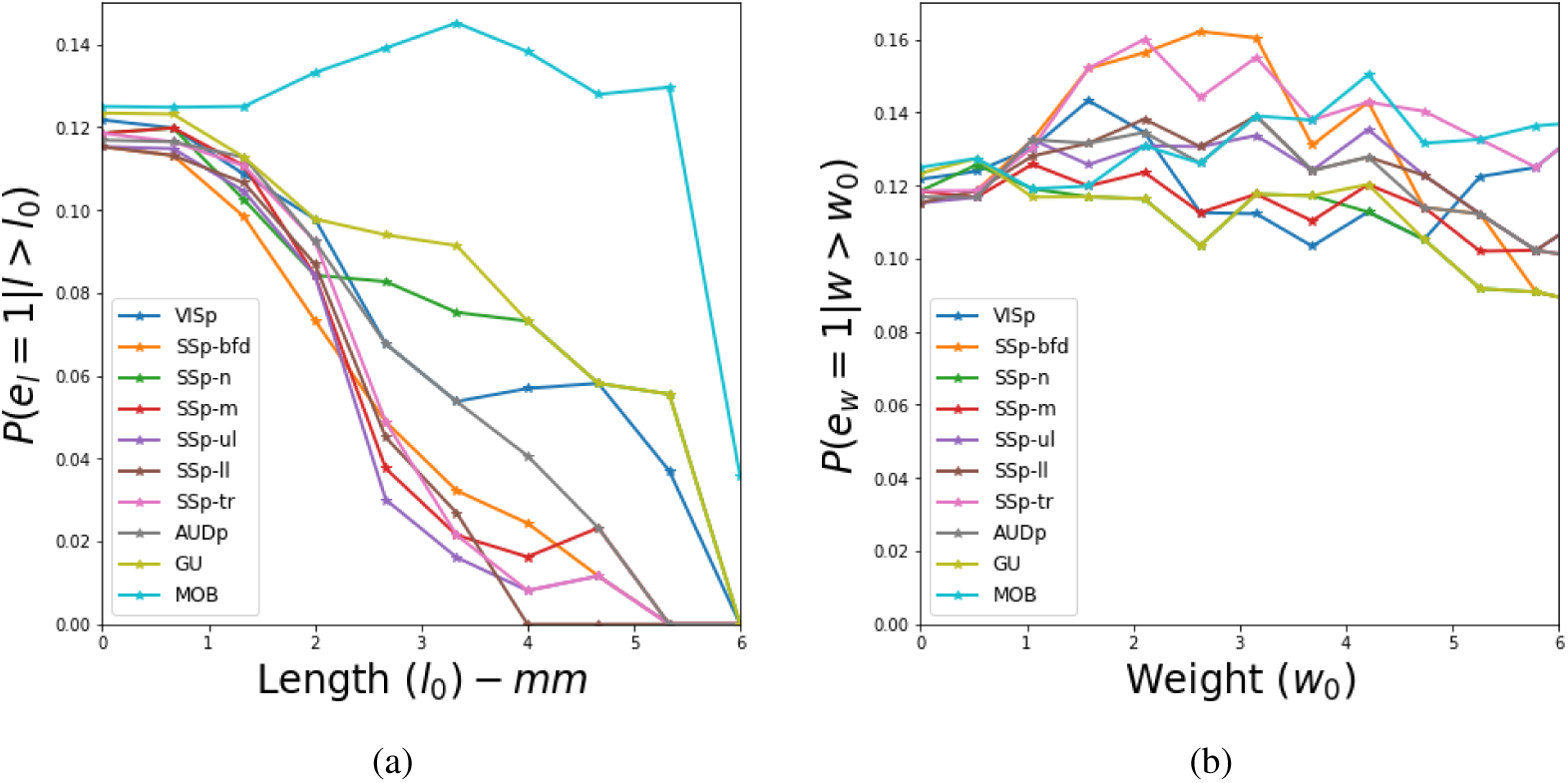
Connections that appear in sensory cascades: (a) Conditional probability that a connection *e_l_* of physical length *l* appears in an activation cascade given that *l > l*_0_. (b) Conditional probability that a connection *e_w_* of weight *w* appears in an activation cascade given that *w > w*_0_.

The right part of Figure 12 shows the corresponding results for the connection weight conditional probability *P*[*e_w_* = 1*|w > w*_0_]. With the exception of the olfactory cascade (MOB), which shows a decreasing trend, the rest of the sensory cascades show a more complex and diverse pattern. The average probability suggests that weight is *not* a significant factor in determining which connections will be part of an activation cascade (perhaps with the exception of edges with very high weights – larger than 8). One reason is that if the weight of a connection is higher than the activation threshold *θ* (which is about one in our modeling results), that edge is sufficient to activate a downstream node, independent of the state of other connections to the same destination node.

These results suggest that *sensory cascades spread in the cortex as a forest fire mostly through short connections connecting physically adjacent regions, rather than through the (relatively few) long connections that connect remote regions*.

### 3.8 Comparison with static network analysis methods

Finally, we asked whether we would we obtain similar results if we had relied on static network analysis metrics and approaches (such as [16, 43, 63]) that do not consider a model of dynamic network activity?

We first ask whether the path centrality metric correlates strongly with more commonly used node centrality metrics, namely: incoming or outgoing strength, betweenness centrality, closeness centrality, pagerank and eigenvector centrality [49]. The results of this comparison are included in the Supp-Info section SI.8. In brief, the conclusion of that comparison is that none of these centrality metrics correlates well with path centrality, computed over all source-target sensory activation paths. The identification of nodes that play an important role in multisensory integration requires an approach that models communication dynamics over the anatomical network – not just the topological properties of the latter [8, 33].

We also examined the similarity between the nine *τ*-core nodes, the five rich-club nodes [29, 18, 54], as well as the thirteen core-periphery nodes computed using Rombach’s method [47]. The complete results appear in SI.8. The overlap between the rich-club and the *τ*-core nodes consists of only the claustrum (CLA) and the supplementary motor region (MOs). The overlap between Rombach’s core nodes and the *τ*-core nodes consists of only MOS and AOB.

In summary, *this analysis highlights the marked differences between ALT-based network diffusion modeling and static network analysis methods based on centrality metrics or core-periphery concepts*.

## 4 Discussion

This study built a suite of tools to quantify and characterize multisensory integration from the perspective of communication dynamics. We chose to analyze the mouse connectome due to the availability of a detailed meso-scale anatomical map and because the problem of multisensory integration is relatively well-studied in rodents [30]. In particular, we analyzed the cortical brain network from the Allen Mouse Connectivity Atlas [42] and focused on “early” dynamics of sensory integration. By *early* we specifically mean the first wave of cortical activity, starting from primary sensory areas and propagating to the whole-hemisphere. This point differentiates our work from earlier models of multisensory integration [67, 46].

The underlying anatomical pathways recruited by diverse sensory modalities (e.g., visual, auditory, somatosensory) branch out rapidly and become increasingly complex as they reach the higher associative cortical areas [35]. To capture the nonlinear and asynchronous nature of these dynamics, we used a diffusion model that can capture both non-linearity and communication delays, called Asynchronus Linear Thresholding (ALT). We found that ALT can closely recapitulate sensory cascades when compared with VSD datasets from sensory stimulation experiments on mice [35].

ALT is a phenomenological model that aims to describe (but not explain mechanistically) the diffusion of information at the ROI level. As such, it does not model the underlying mechanisms of neuronal communication through chemical or electrical synapses, and it is also quite different than neural mass models that capture coarse grained activity of large populations of neurons and synapses [9]. Similar to neuroimaging modalities of similar spatial resolution, such as task-based fMRI, ALT aims to capture how a certain activity, namely the stimulation of a primary sensory region, causes the activation of other brain regions. A difference with fMRI or MEG however, is that the resulting activation cascades can be analyzed to infer interactions between ROIs that participate in the cascade, as described above. Additionally, ALT depends on the communication delays and weights of the connections between brain regions. Thus, it produces a timeline of activation events, one for each ROI that participates in the cascade. The length of the time interval between activation events, at least in relative terms, can be compared with experimental results from neuroimaging modalities that have fine temporal resolution, such as VSD (as in this study) or calcium imaging.

On the other hand, the ALT model cannot capture more complex dynamics, such as sustained oscillations at certain frequencies or feedback from a newly activated region back to regions that were activated earlier. Modeling such dynamics would require more elaborate neural mass models – their validation however would require whole-brain neuroimaging data of fine spatial and temporal resolution (higher temporal resolution than fMRI and higher spatial resolution that EEG). Additionally, those more complex models may not be necessary when the goal is to map the feedforward initial propagation of brain activity after a sensory stimulation.

Our primary result is that relatively few cortical regions are responsible for integrating almost all sensory information. This finding supports the idea that multisensory integration is performed through an “hourglass architecture”. The benefit of an hourglass architecture is that it first reduces the input dimensionality of the sensory stimulus at few intermediate-level modules that reside at the “hourglass waist”. Second, it re-uses those compressed intermediate-level representations across higher-level tasks, reducing redundancy between the latter. The hourglass analysis framework was first developed in [50] and it has been recently applied in the connectome of *C.elegans* [51]. There are two fundamental differences in our study: i) we rely on communication dynamics while [51] uses a predefined set of “routing mechanisms” to construct ensembles of sensory pathways, and ii) the sources and targets of the *C.elegans* cascade are the sensory and motor neurons, respectively. Nonetheless, it is interesting that the “hourglass architecture” emerges in both studies. One possibility is that this architecture is selected by evolution because it drives a network towards reusing a small set of intermediate functions in constructing a range of redundant output functions.

Rather than studying each sensory cascade in isolation, our analysis framework is based on the combination of all sensory cascades. Comparative and competitive cascades have been studied in [34] to quantify the combined effect of multiple cascades, using simultaneously activated source nodes. Sensory stimuli of different modalities, however, do not need to arrive at the cortex simultaneously in order to be integrated [41]. Different sensory stimuli travel at different speeds through body receptors [68]. The analysis framework that we followed constructs uni-sensory cascades and merges their activation paths. We have also experimented with cascades originating from two simultaneously activated sources (see Section SI.7). The hourglass architecture is still observed in that case, and the core nodes are mostly the same.

There is mounting evidence that points to the fact that many computations in the brain are multi-sensory. Even primary sensory cortices respond to sensory stimuli from different modalities. For instance, there is significant activity in olfactory cortex after delivering taste stimuli to the tongue [25]. Or, many units in the auditory cortex respond to visual stimuli [37]). A number of studies have shown primary-like responses from multiple sensory modalities [5, 11], advancing the notion that much of cortex is multi-sensory [13]. Our results shed new light on this debate, and support the findings that many regions in early sensory cascades are multi-sensory, the most important of which are the claustrum and the posterior parietal cortex.

The claustrum is known for its anatomical uniqueness [1] and its precise function has been enigmatic [64, 28]. The late F. Crick has hypothesized that the claustrum may be a potential gateway to consciousness [7]. More recent results have demonstrated that the claustrum is important in gating sensory information, and in attention mechanisms in visual perception [64]. These and other studies provide strong evidence that the claustrum is a crucial node for multisensory integration. Our findings are in agreement with this growing body of work.

The posterior parietal cortex is the second most important region for MSI, according to our analysis. This stems from its strong and immediate connectivity to primary sensory regions and its projections to motor areas. PTLp’s connectivity points to its role in integrating sensory information to direct immediate motor commands [65]. The experiments of Nikbakht et al. [40] provide direct evidence that PTLp is multisensory both at the behavioural and neurophysiological levels and it provides sensory independent information about the orientation and categorization of objects in the environment. We refer the reader to [36] for further discussion.

Three additional core nodes have been associated with specific sensory modalities in the past: Ventral auditory area (AUDv), Supplemental somatosensory area (SSs), and Lateral visual area (VISl). However, all of them have also been implicated to some extent with multisensory processing. For instance, Hishida et al. [21] found that activity propagating to the parietal association area passes through the ventral auditory region, irrespective of the sensory stream source (visual, auditory or somatosensory). Similarly, [31] suggests that SSs has a major role in bringing context to the sensory pathways [48]. VISl is on the dorsal visual stream and is associated mostly with spatial location and action guidance [27], while the other secondary visual areas participate in the ventral stream [20].

An important next step is the inclusion of subcortical regions in the diffusion model. Various nuclei in the thalamus or the superior colliculus are known to be crucial in MSI [58]. An integrated model of both cortical and subcortical activity, in the context of sensory integration, may help to explain the complex cortico-thalamic feedback mechanisms [22, 62]. Disentangling such communication dynamics requires working at finer spatial resolutions, and including distinct cortical layers (laminar divisions) [14]. Incorporating high-resolution [24] and cell type specific [19] information about connectivity in our pipeline is an interesting direction, that can allow segregation of different communication channels. By increasing the spatial resolution and the complexity of the model dynamics, we could capture more complex population-level activity patterns (e.g., Wilson-Cowen [9]) and better understand the role of mesoscale network topology in constructing a coherent perceptual state from raw sensory streams.

## Supplementary Information

### SI.1 Sources of sensory cascades

The ten cortical regions we consider as sources of sensory activation cascades are shown in Table SI-1.

**Table SI-1:** Sources of sensory cascades

| Acronym | Description | Major region | Size (number of voxels) |
| --- | --- | --- | --- |
| AUDp | Primary auditory area | Isocortex | 2689 |
| VISp | Primary visual area | Isocortex | 6227 |
| GU | Gustatory areas | Isocortex | 2104 |
| SSp-n | Primary somatosensory area, nose | Isocortex | 1358 |
| SSp-bfd | Primary somatosensory area, barrel field | Isocortex | 10306 |
| SSp-ll | Primary somatosensory area, lower limb | Isocortex | 3254 |
| SSp-m | Primary somatosensory area, mouth | Isocortex | 2924 |
| SSp-tr | Primary somatosensory area, trunk | Isocortex | 4799 |
| SSp-ul | Primary somatosensory area, upper limb | Isocortex | 5406 |
| MOB | Main olfactory bulb | Olfactory Areas | 16978 |

### SI.2 Connectome filtering

**Figure SI-1:**
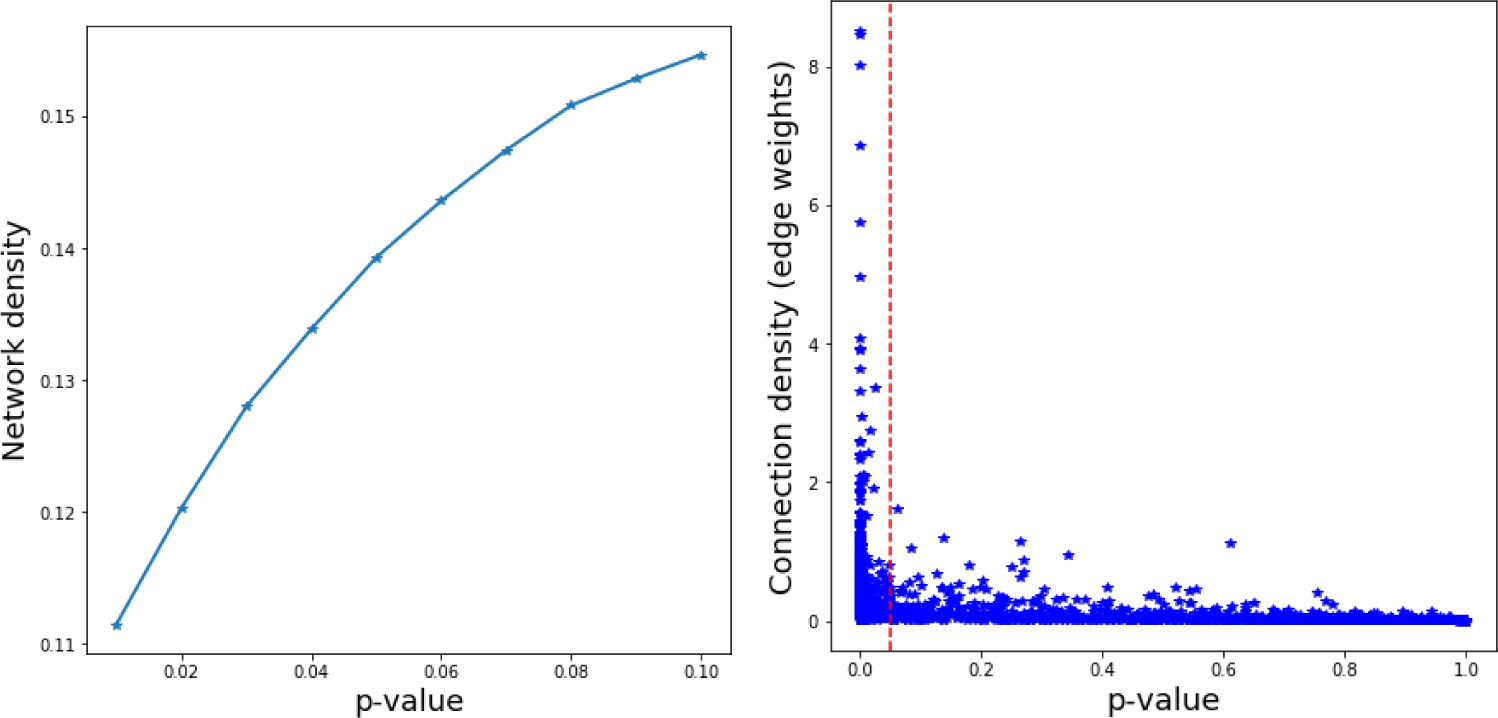
Connectome filtering. **Left:** Network density versus edge p-value. **Right:** Connection density (edge weight) versus edge p-value

We have repeated the analysis for the following p-values: 0.01, 0.02*, · · ·* 0.1. When the p-value is smaller than 0.03, the network becomes disconnected. When the p-value is higher than 0.04, the core nodes remain the same as in Table 8d.

Figure SI-1 shows the network density for different p-values (left) as well as the connection density (edge weight) versus p-value for each structural connection among the 67 ROIs we consider (right).

### SI.3 The activation cascade of each sensory source

The ten activation cascades, one for each sensory source, are shown in Figures SI-2 through SI-11.

### SI.4 Sensitivity to activation threshold *θ*

Recall that if *θ* is higher than 0.98, the cascade of some sources will not be complete. We have also computed the core nodes for two lower values of *θ*: 0.9 and 0.95. The 70%-core remains exactly the same. The 90%-core for *θ* = 0.9 includes PERI instead of ECT.

### SI.5 Analysis of “disagreement cases” between VSD data and ALT modeling results

In this section we analyze the cases in which the VSD experimental results predict a different temporal ordering than the ALT modeling results. Recall that a disagreement refers to a pair of ROIs (e.g., X and Y) for which the activation order in ALT is different than that in VSD. Thus, every disagreement case involves two different ROIs.

We first split the ROI pairs in two sets, the disagreement cases and the agreement cases. We measured the Euclidean distance between the center of the two ROIs in each pair of the two sets. The hypothesis that the sample mean of these distances is the same could not be rejected with a p-value of 5%.

We did the same for the connection weight between connected ROI pairs, comparing the average weight of pairs in the set of agreements and the set of disagreements. Again, the hypothesis that the sample mean of these distances is the same could not be rejected with a p-value of 5%.

So, the distance or connection weight between two ROIs does not predict whether they will be a disagreement case.

Are there certain ROIs that appear in surprisingly many disagreement cases? To answer this question, we measure the number of times *X* that each ROI appears in the *N* disagreement cases that are observed in the cascade of a certain sensory modality (considering the datasets form all five animals). If an ROI appears in *k > µ* + 3*σ* disagreement cases, we conclude that it is significantly over-represented in the group of disagreement cases. The mean and standard deviation in the previous inequality are calculated based on the null model that the disagreements involve randomly chosen pairs of distinct ROIs. So, if *K* is the number of ROIs and we have *N* disagreements (for a specific sensory cascade), then the probability of selecting a specific ROI in each pair is:

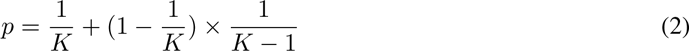

Given that we sample N pairs, the null model is that we will sample each ROI a number of times *X*, where *X* follows the *Binomial*(*N, p*). So, *µ* = *N p* and *σ*^2^ = *N p* (1 *− p*).

Figure SI-12 shows the results for this study. Note that in each of the five cascades, it is only one or two ROIs that are significantly over-represented in disagreement cases. In the visual cascade for instance, it is the SSp-bfd ROI that is present in surprisingly many disagreements.

We further analyzed these disagreements in which one of the two ROIs is an outlier, asking whether the second ROI is also over-represented. Figure SI-12f identifies such ROI pairs – four of the cascades have only one pair while the Forelimb stimulation cascade has none. ACAd is an ROI that appears in many disagreements in the visual, whisker and tone stimulation cascades – note that this ROI appears at the boundary of the cortical surface at the VSD datasets and it is only partially visible. So, it is likely that the VSD data do not reflect accurately the time at which that region is activated after each stimulation. To a smaller degree, the same may be true for TEa, which is the outlier in the hindlimb stimulation cascade.

### SI.6 Similarity between activation cascades with single and complete linkage

Figure SI-13 shows hierarchical clustering dendrograms quantifying the similarity between the ten activation cascades based on single (left) and complete (right) linkage. The corresponding average linkage plot is shown in Figure 5-b.

### SI.7 Activation cascades when two sensory sources are activated simultaneously

We have also considered the case of two simultaneously active sources, considering all possible pairs of sources (10×9/2=45 cascades). The hourglass analysis is summarized in Figure SI-14. The core nodes are the same with the single-source case, except that the anterior cingulate area – dorsal part (ACAd) and the Ectorhinal area (ECT) are replaced by the perirhinal area (PERI).

### SI.8 Comparison with other network analysis metrics

We first ask whether the path centrality metric correlates strongly with more commonly used node centrality metrics, namely: incoming or outgoing strength (the equivalent of “degree” for directed and weighted networks), betweenness centrality (fraction of all shortest paths traversing a node), closeness centrality (inversely related to average shortest path distance from that node to any other node), pagerank and eigenvector centrality (two related “influence” metrics that assign a higher score to a node that is connected to other highly-scored nodes compared to a node that has the same number of connections to low-scored nodes) [49]. Given that we are mostly interested in the dissemination of sensory information over the network, one may expect that the pagerank and eigenvector centrality metrics would be more highly correlated with path centrality because such cascades do not necessarily follow shortest paths [44].

**Table SI-2:** Top-five nodes according to different centrality metrics

| Centrality metric | Top-five nodes |
| --- | --- |
| Out-strength | ENTl - CA2 - PIR - FRP - EPv |
| In-strength | CA1 - MOs - ENTl - DP - FRP |
| Betweenness | ENTl - CLA - PERI - VISI - MOs |
| Closeness | ENTl - PERI - CLA - PIR - VISI |
| Pagerank | VISI - RSPagl - VISpm - FRP - RSPv |
| Eigenvector | MOs - FRP - AOB - CLA - MOp |
| **Path Centrality** | CLA - SSs - PTLp - AUDv - MOs |

Table SI-2 shows the top-five nodes in the network for each centrality metric. Figure SI-15 shows scatter plots for the previous centrality metrics, comparing each of them with path centrality. The plots also show Kendall’s *τ* rank correlation coefficient. All centrality metrics are computed using ***networkx*** [17].

Strength (either for outgoing or incoming edges) only considers the local connections of each node; nodes in the hippocampal formation (CA1, CA2, ENTl) are among the most strongly connected, while none of the *τ*-core nodes ranks highly in terms of strength. The betweenness and closeness metrics are both based on shortest paths; the lateral entorhinal (ENTl) ranks highest in terms of that metric, while the claustrum (CLA) is the only *τ*-core node in the top-5 according to these two metrics. The highest ranked node based on pagerank is the lateral visual area (VISl) while the highest ranked node based on eigenvector centrality is (by far) the secondary motor area (MOs).

**Table SI-3:**
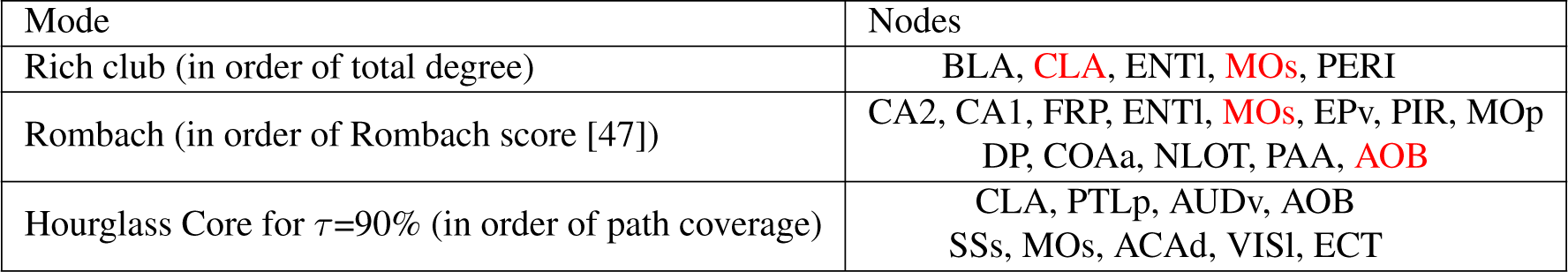
Rich-club nodes, Rombach core nodes nodes, and hourglass core nodes (*τ* =90%): the overlap between the first two sets with the hourglass core nodes is highlighted in red.

| Mode | Nodes |
| --- | --- |
| Rich club (in order of total degree) | BLA, **CLA**, ENTl, **MOs**, PERI |
| Rombach (in order of Rombach score [47]) | CA2, CA1, FRP, ENTl, **MOs**, EPv, PIR, MOp<br>DP, COAa, NLOT, PAA, **AOB** |
| Hourglass Core for $\tau=90\%$ (in order of path coverage) | CLA, PTLp, AUDv, AOB<br>SSs, MOs, ACAd, VISI, ECT |

Table SI-3 shows the five rich-club nodes. The rich-club analysis is performed on unweighted and undirected networks and the rich-club coefficient is computed based on 1000 random surrogate networks, as described in [29]. The rich-club coefficient peaks at 1.15 for nodes with total degree over 23. Only two of the nodes in the rich-club overlap with the hourglass *τ*-core (CLA and MOs).

We also performed a core-periphery analysis on weighted but undirected networks using Rombach’s method [47]. The core nodes according to this method are reported in Table SI-3. Again, only two of the nodes in that core set overlap with the hourglass *τ*-core (MOs and AOB).

**Figure SI-2:**
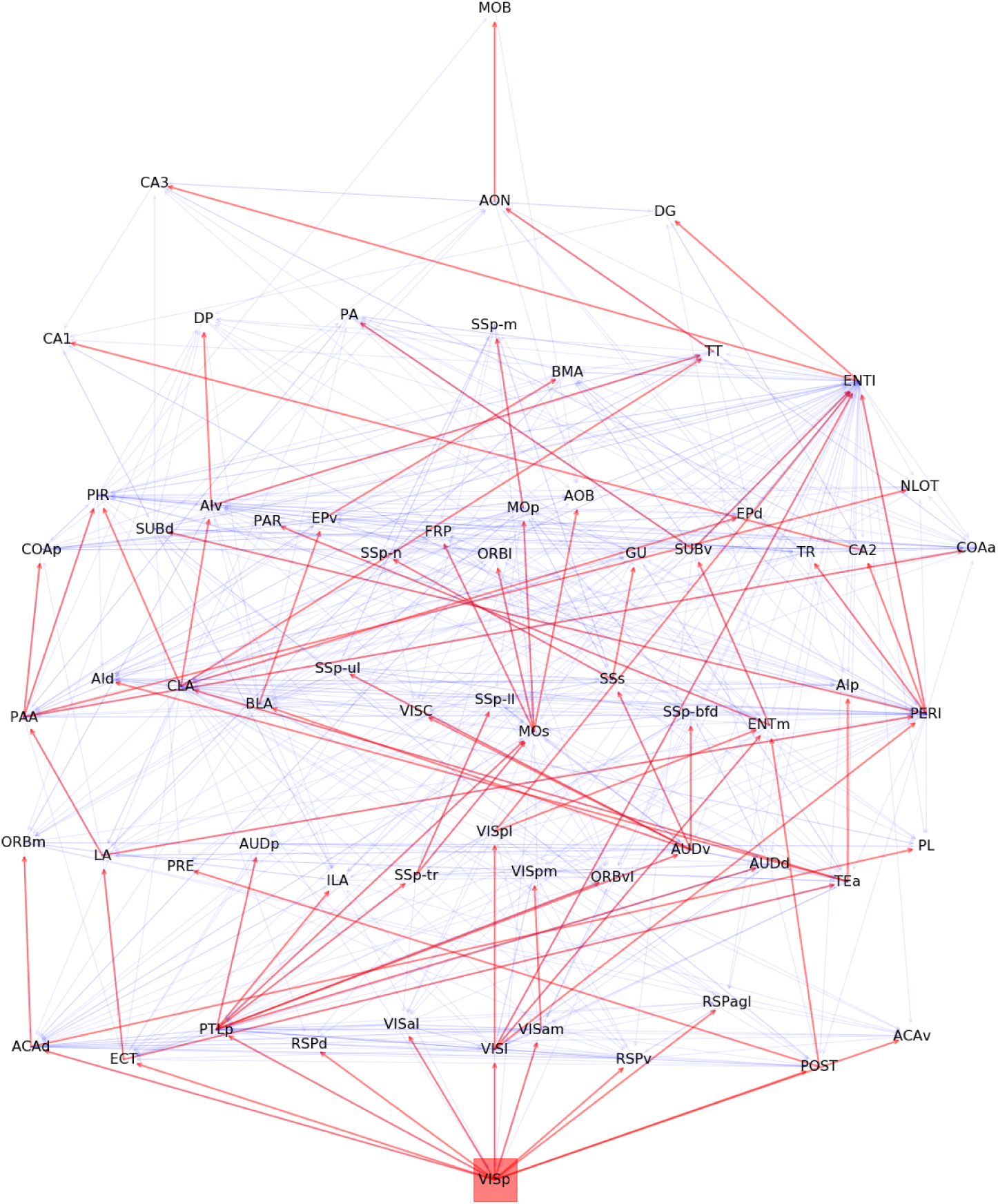
Visual cascade. (source: VISp)

**Figure SI-3:**
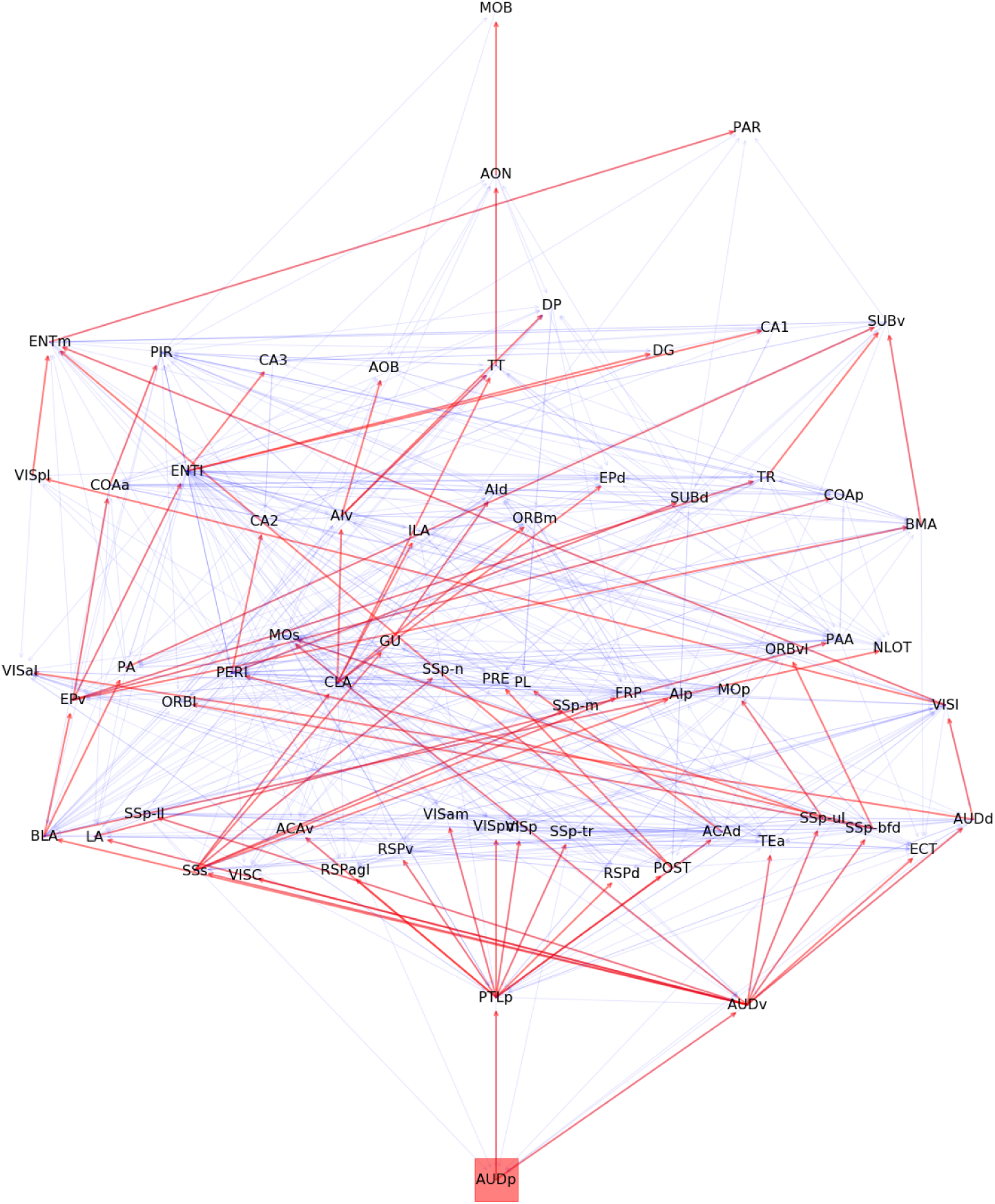
Auditory cascade. (source: AUDp)

**Figure SI-4:**
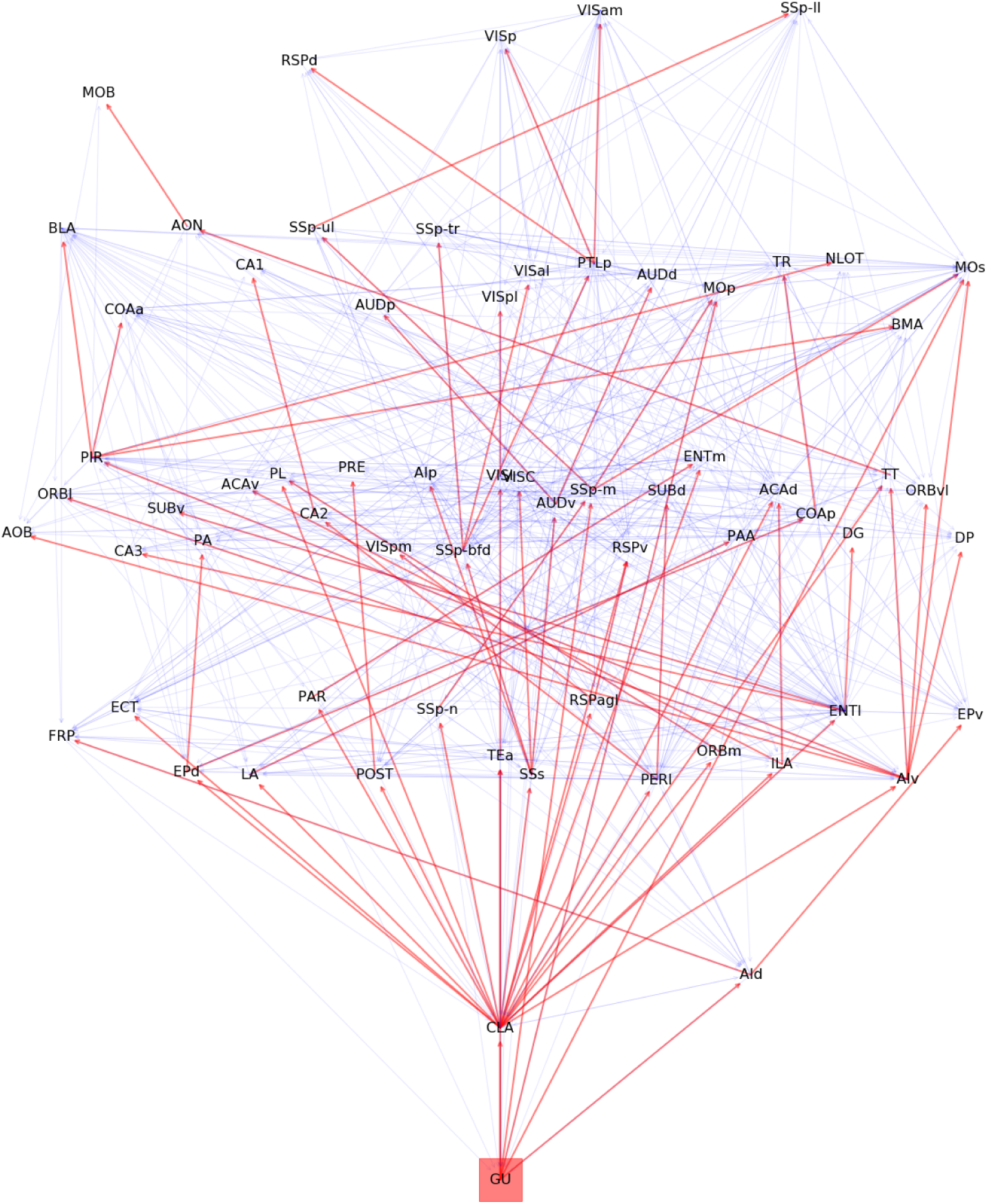
Gustatory cascade. (source: GU)

**Figure SI-5:**
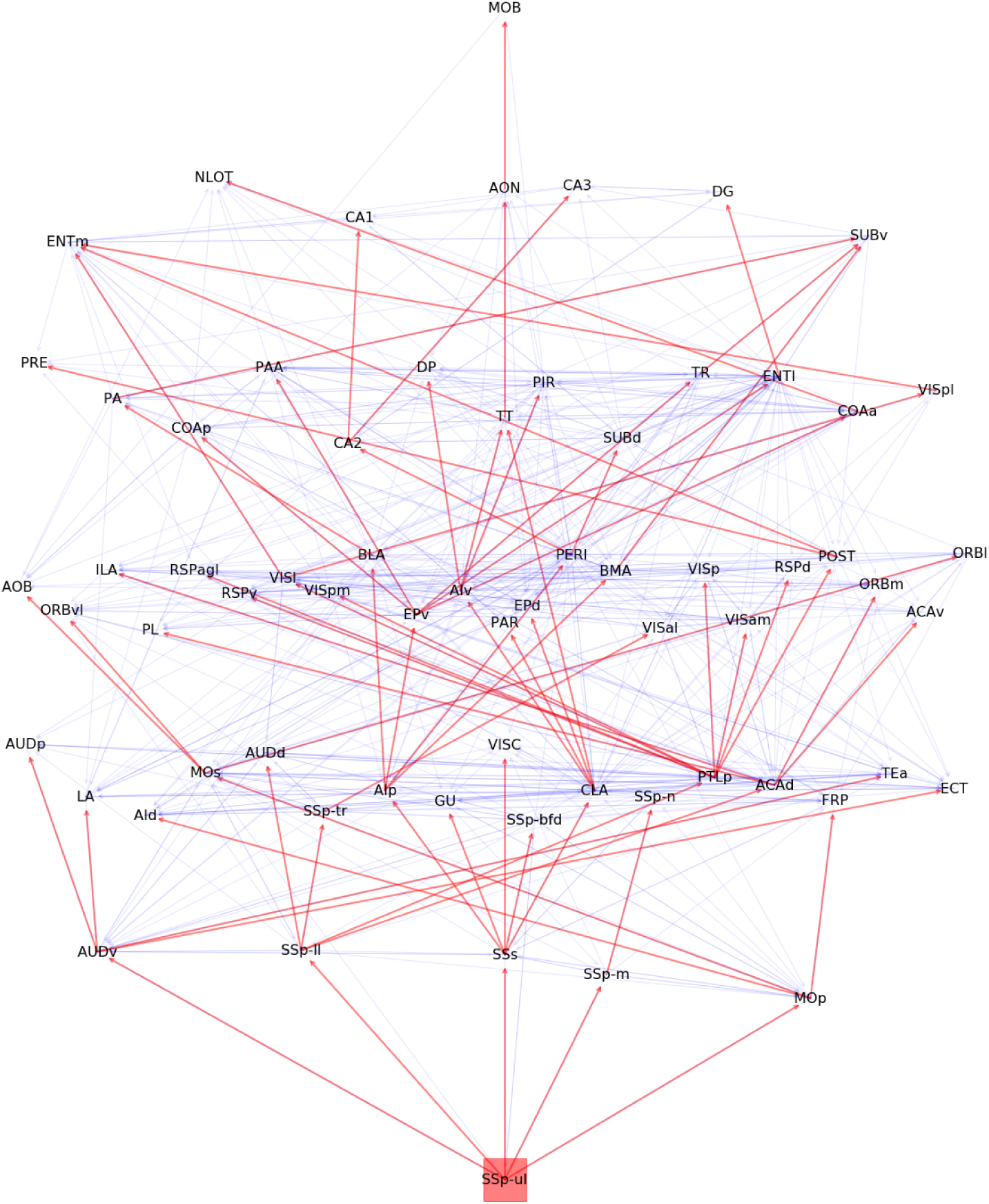
Upper-limb somatosensory cascade. (source: SSp-ul)

**Figure SI-6:**
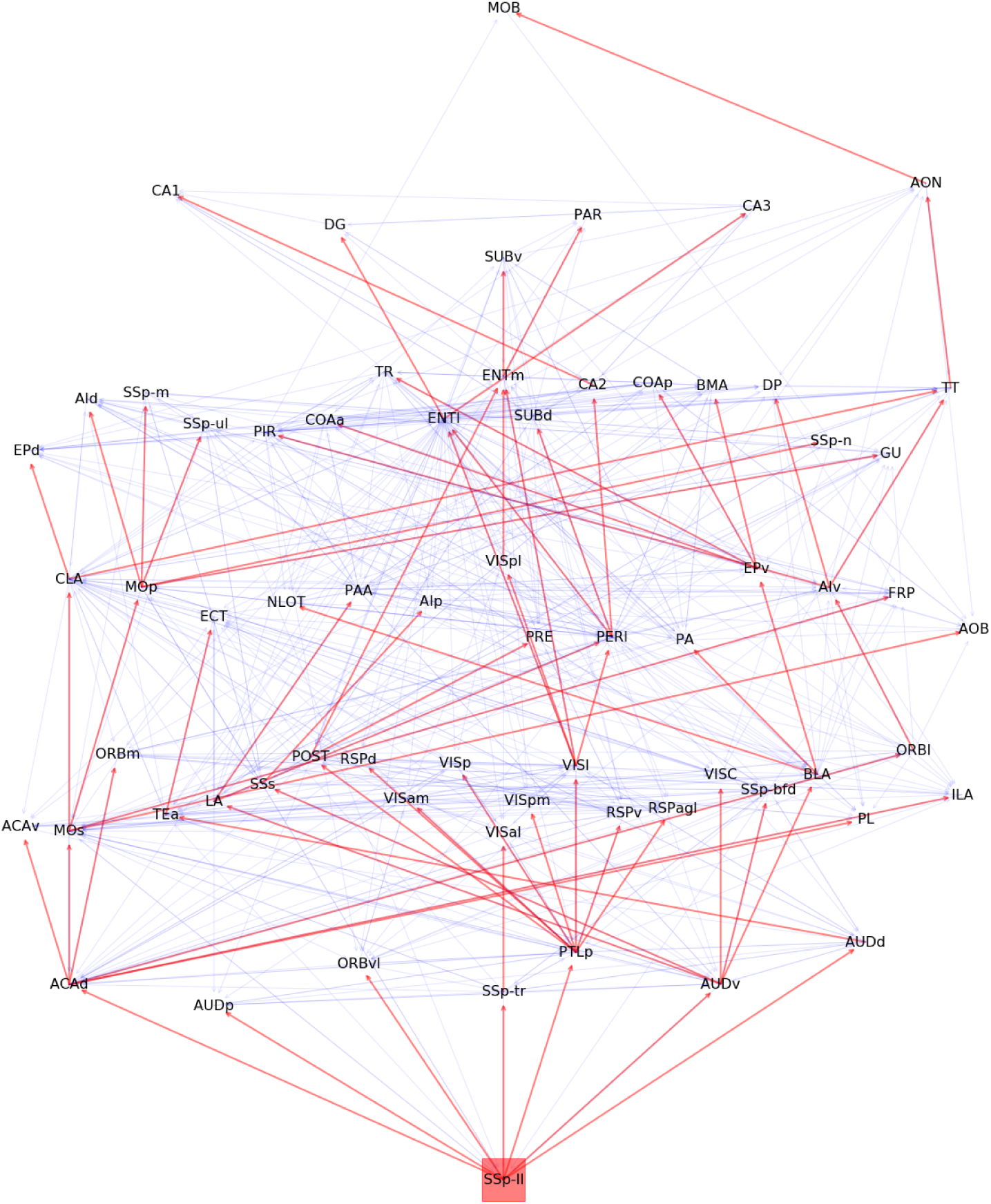
Lower-limb somatosensory cascade. (source: SSp-ll)

**Figure SI-7:**
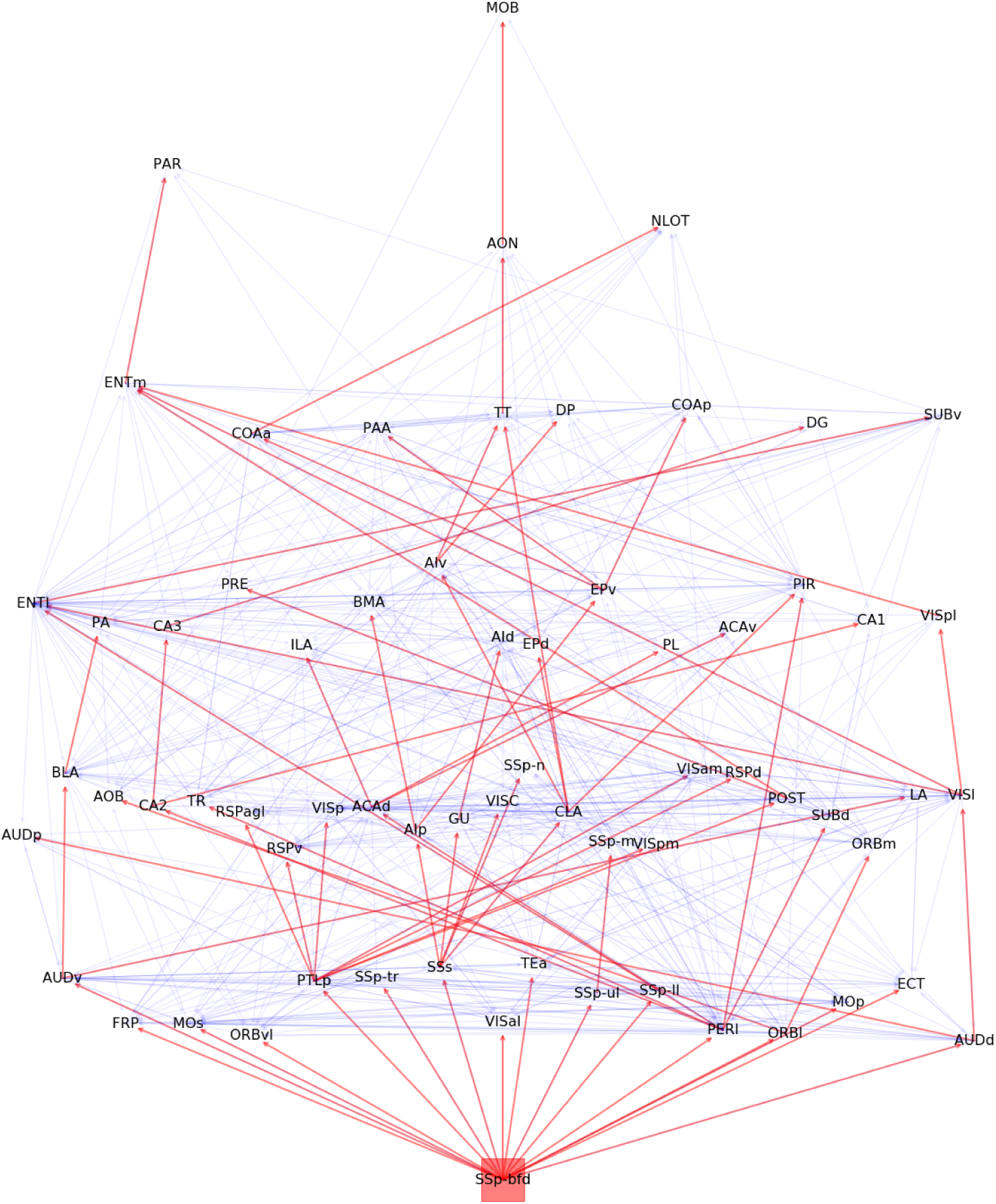
Whiskers somatosensory cascade. (source: SSp-bfd)

**Figure SI-8:**
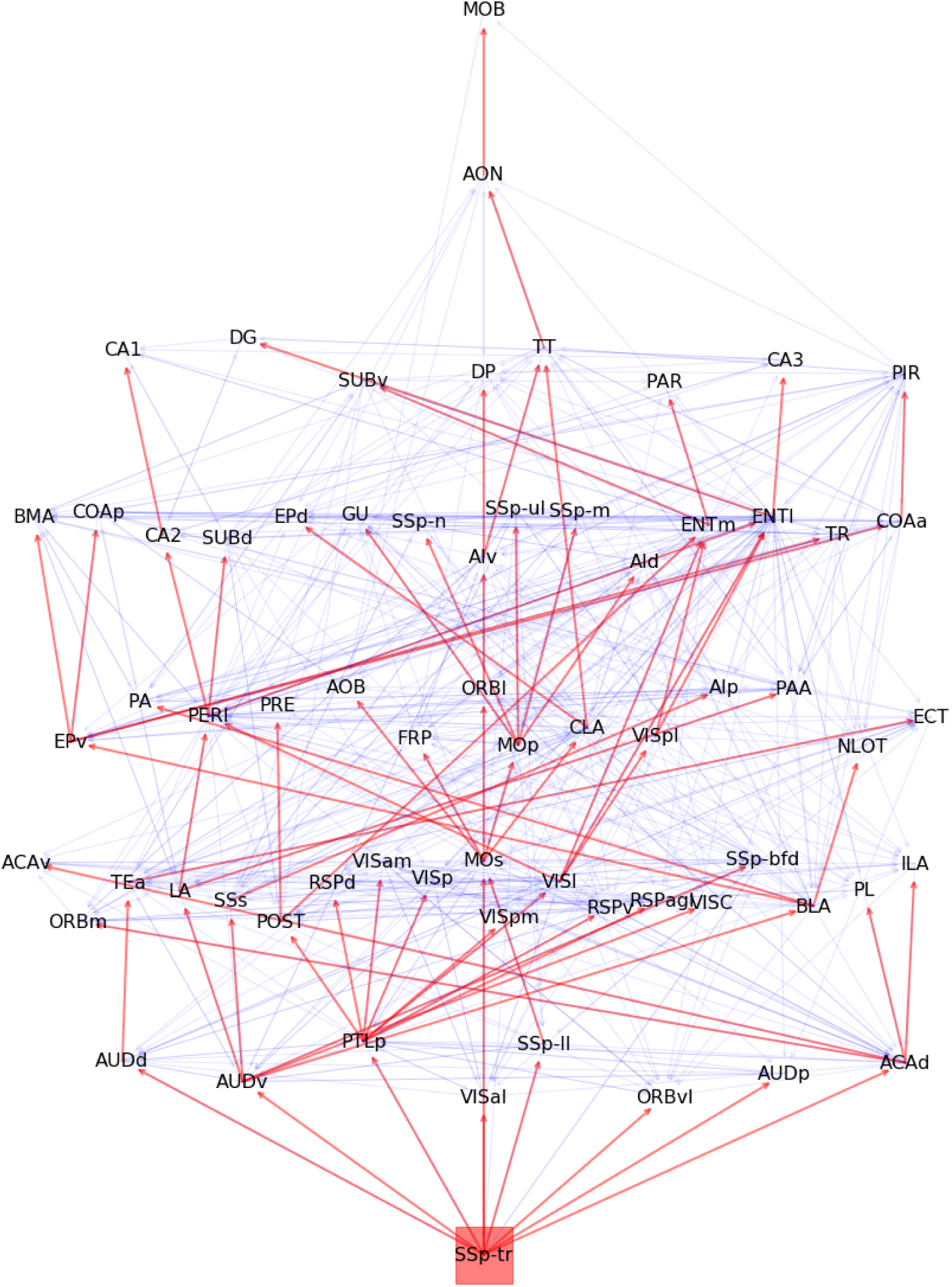
Trunk somatosensory cascade. (source: SSp-tr)

**Figure SI-9:**
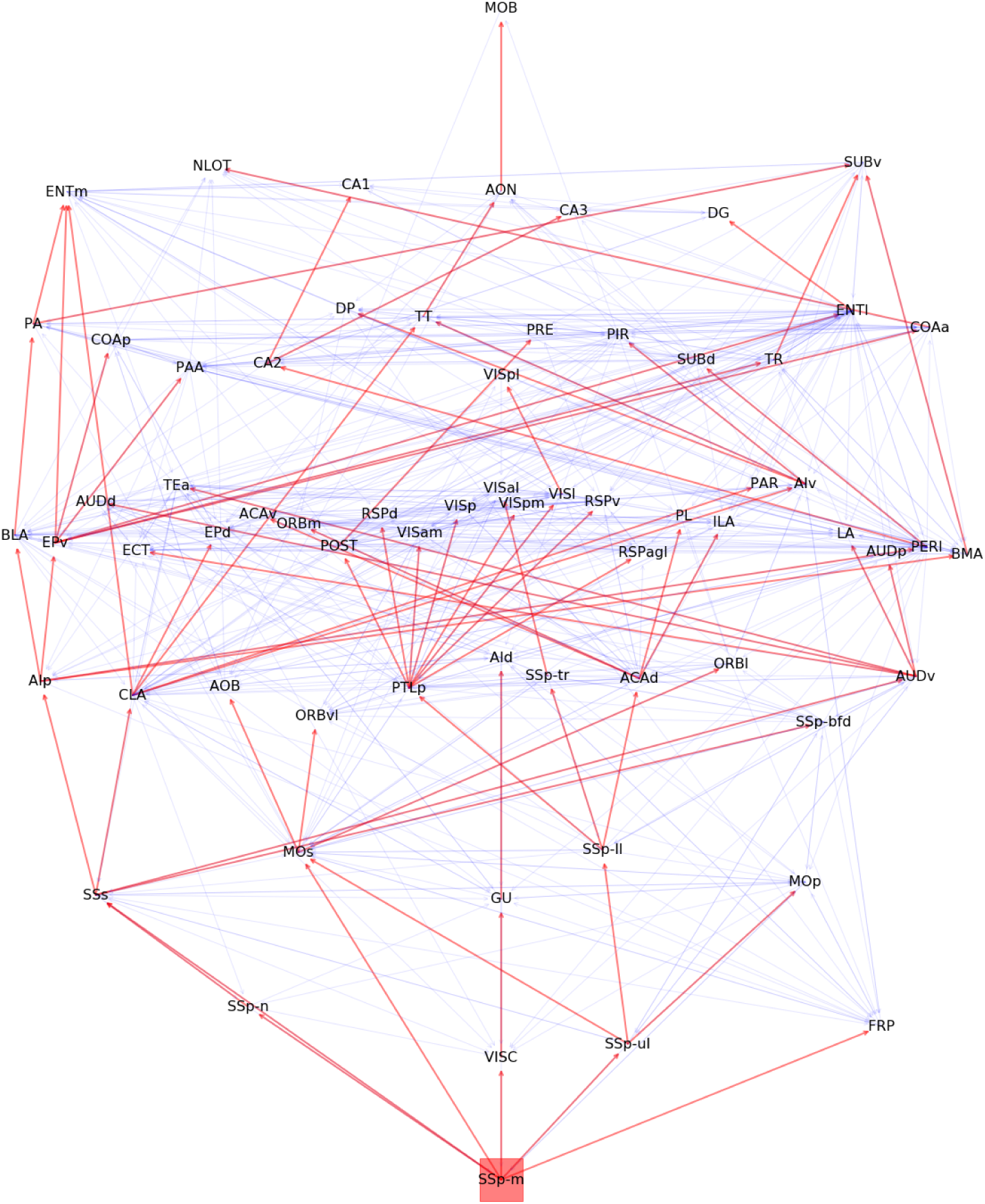
Mouth somatosensory cascade. (source: SSp-m)

**Figure SI-10:**
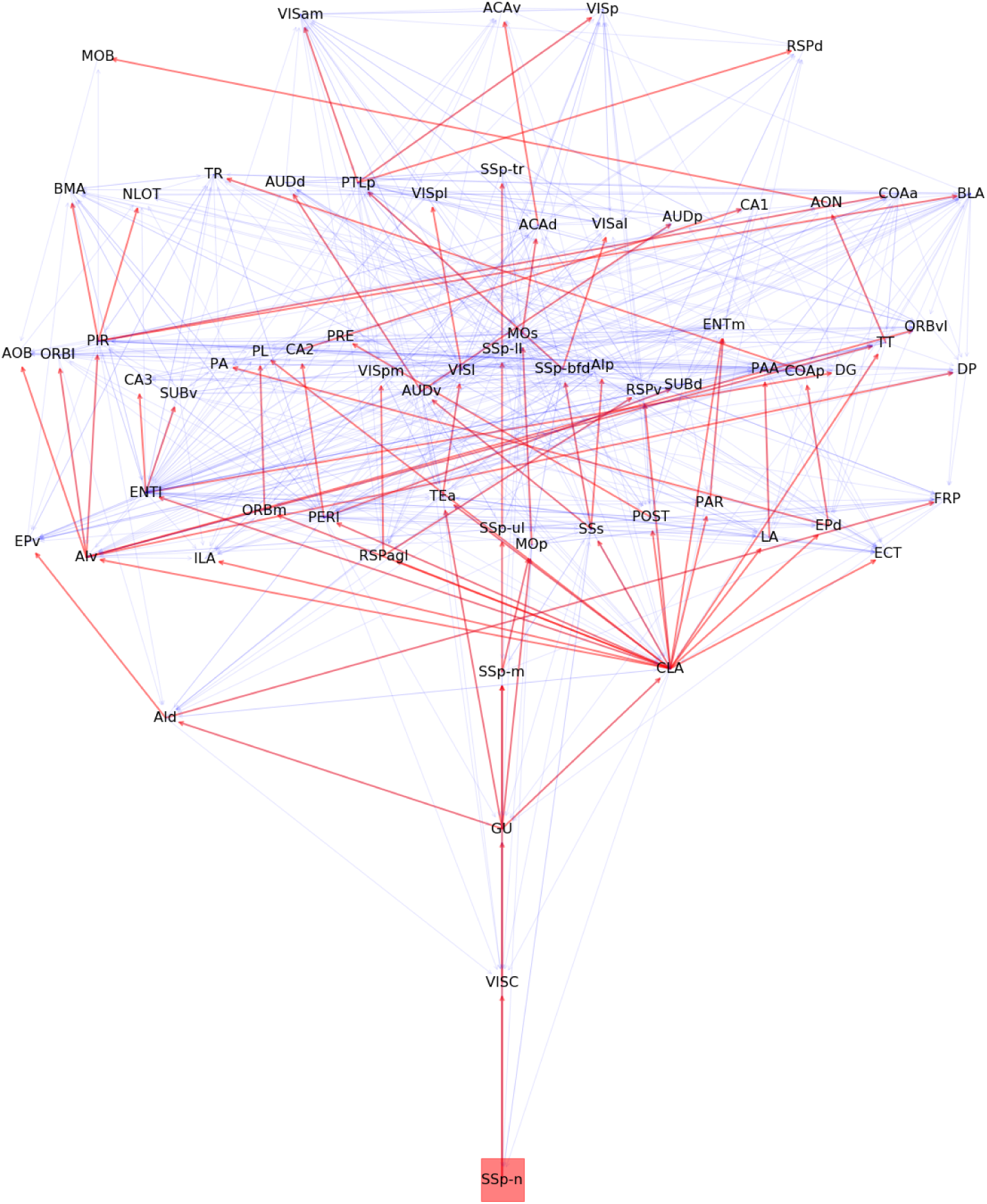
Nose somatosensory cascade. (source:SSp-n)

**Figure SI-11:**
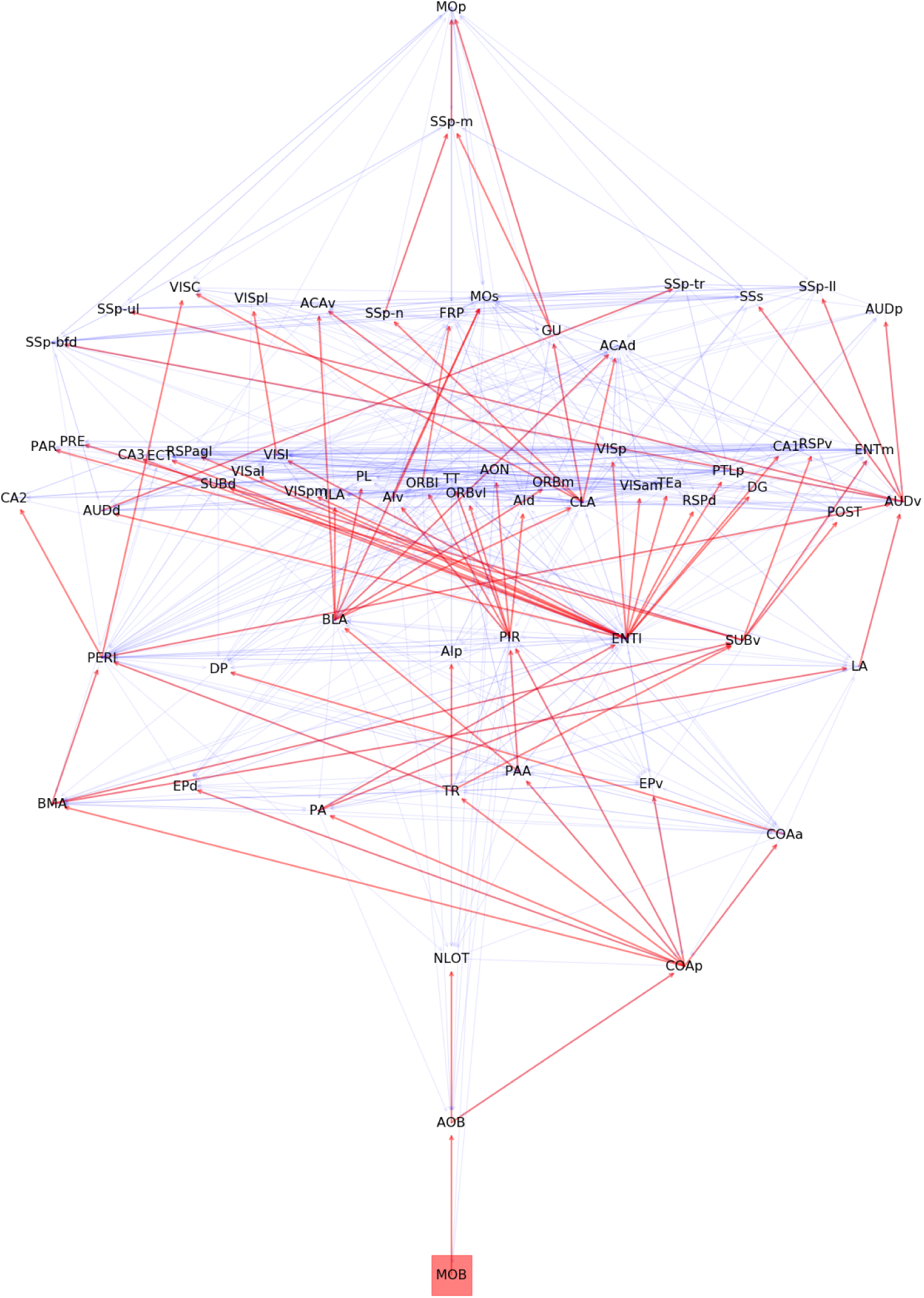
Olfactory cascade. (source: MOB)

**Figure SI-12:**
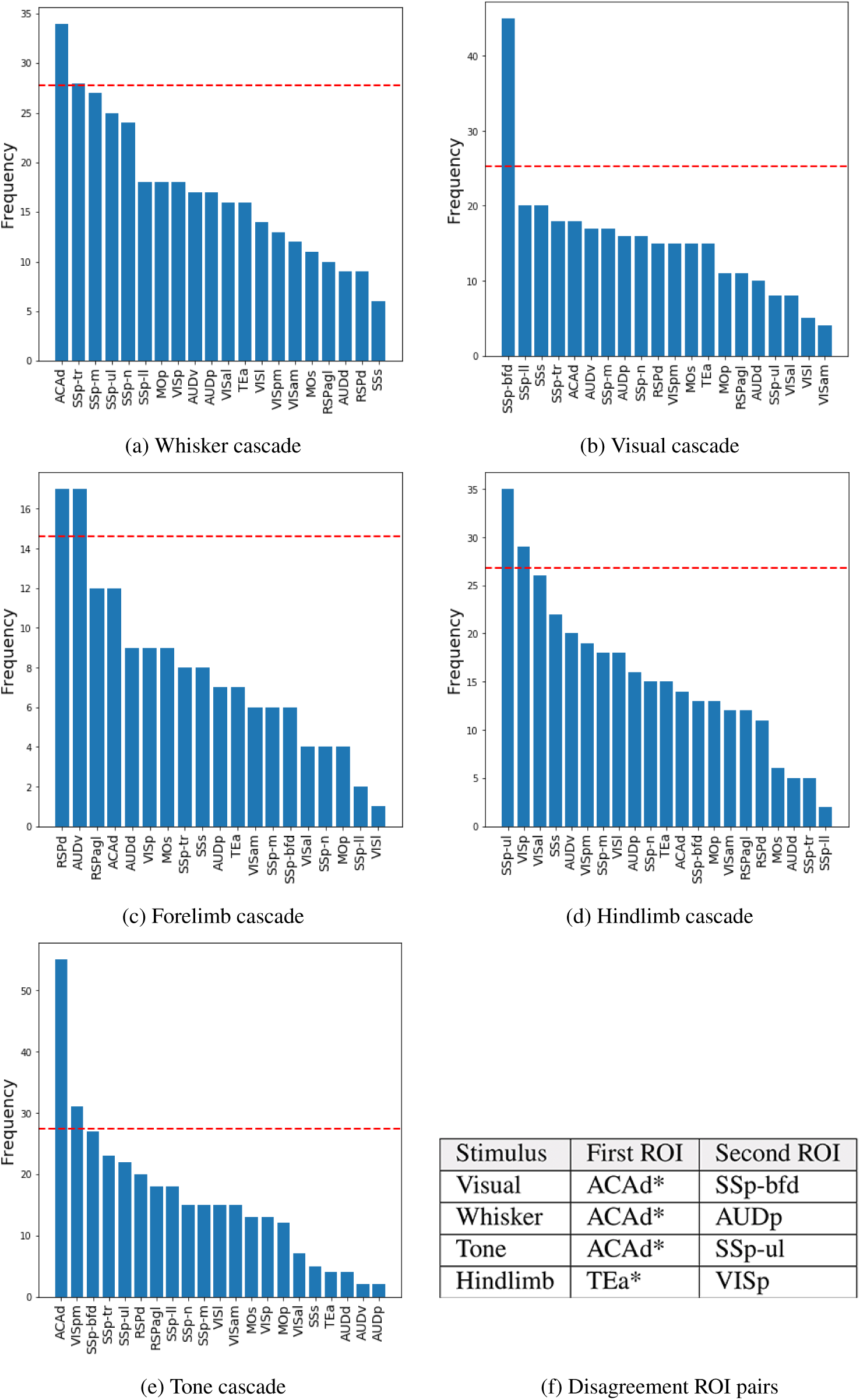
(a)-(e): The number of disagreements between VSD and ALT that involve each ROI, over the 21 ROIs that appear in the VSD data. The outlier ROIs have frequencies that exceed the red dashed line. (f) Disagreement ROI pairs that appear in all five animal datasets - each row corresponds to a different stimulus. ROIs at the boundary of the VSD visible cortical surface are marked by a star.

**Figure SI-13:**
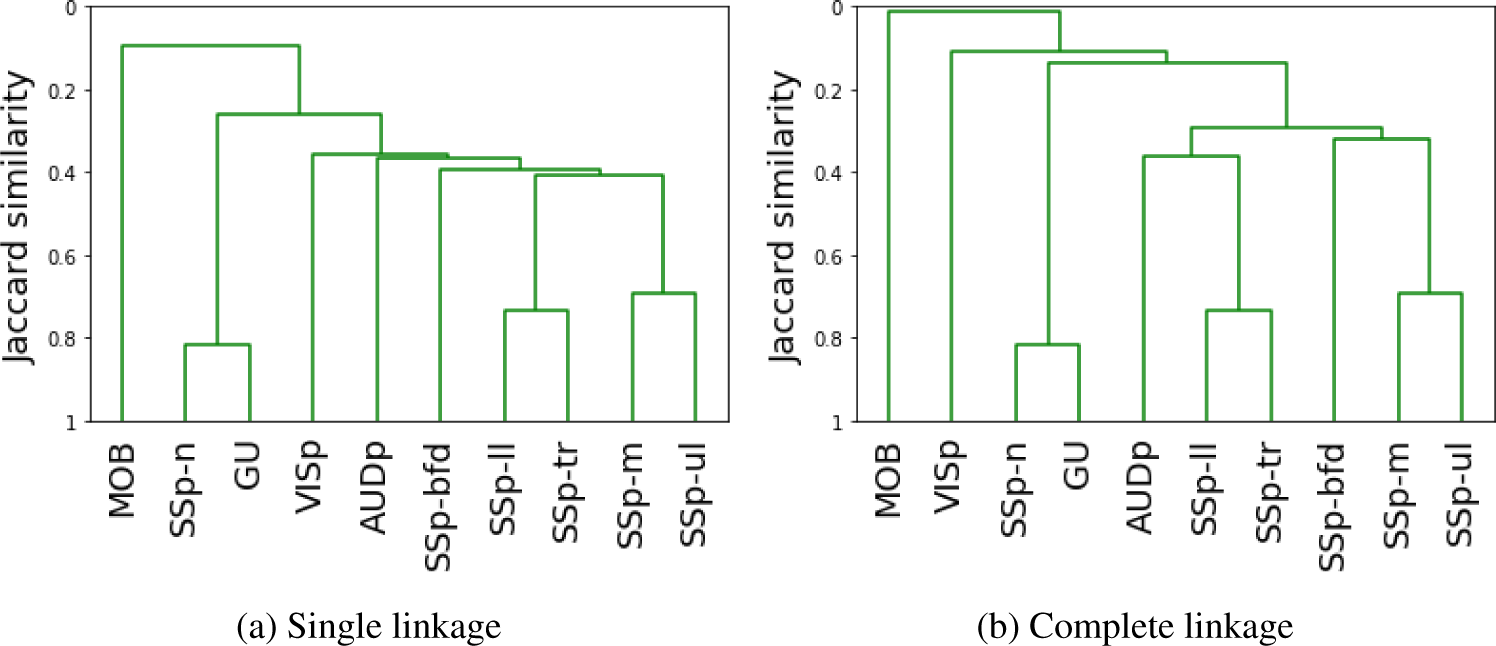
Similarity between activation cascades.

**Figure SI-14:**
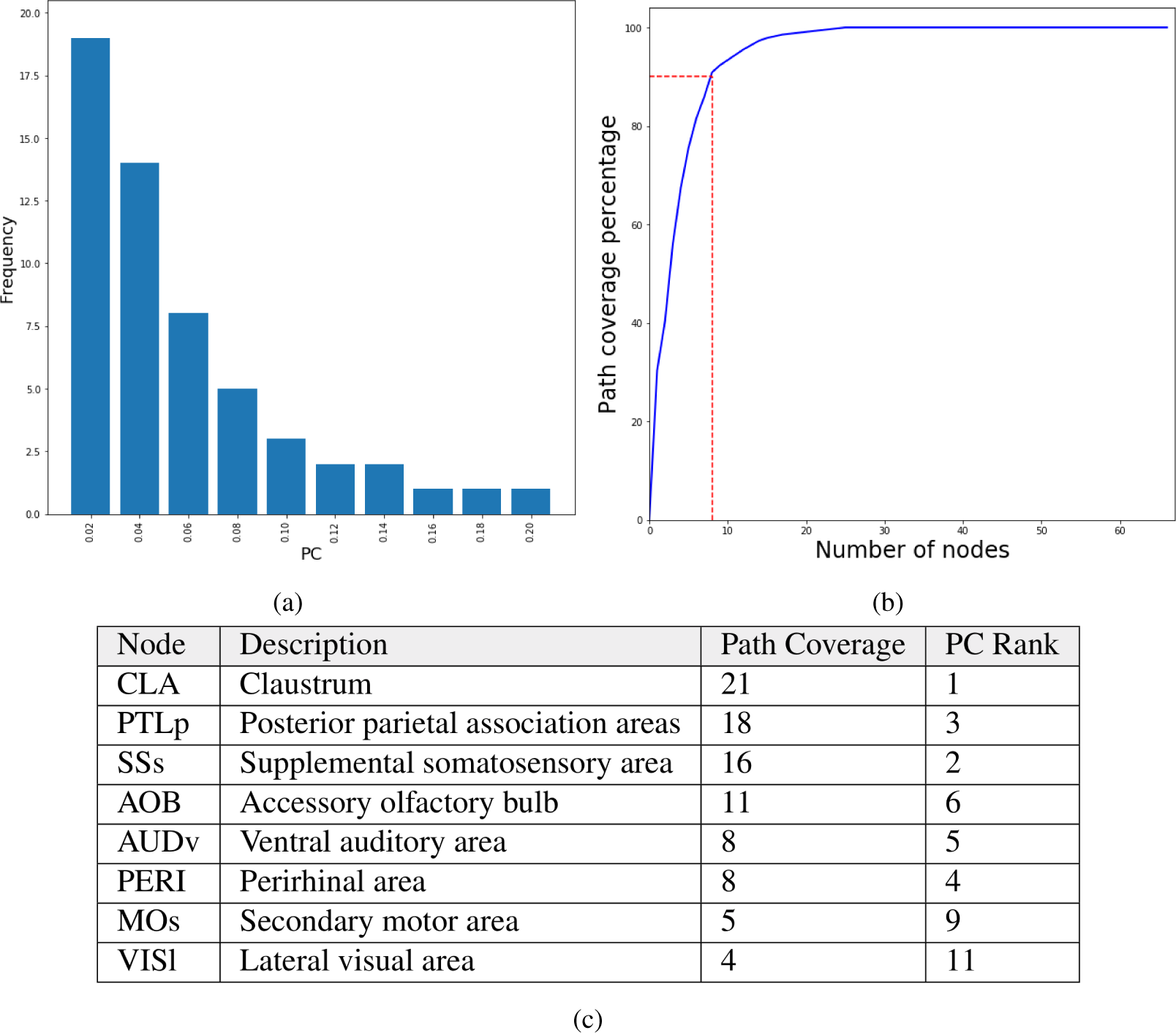
Simultaneous activation of two source nodes: (a) Path Centrality (PC) histogram for the 67 cortical nodes, considering all source-target paths across the 10×9/2=45 activation cascades. (b) Cumulative path coverage by the top-X core nodes for X=1···67. (c) Eight nodes are enough to cover *τ* = 90% of all paths.

**Figure SI-15:**
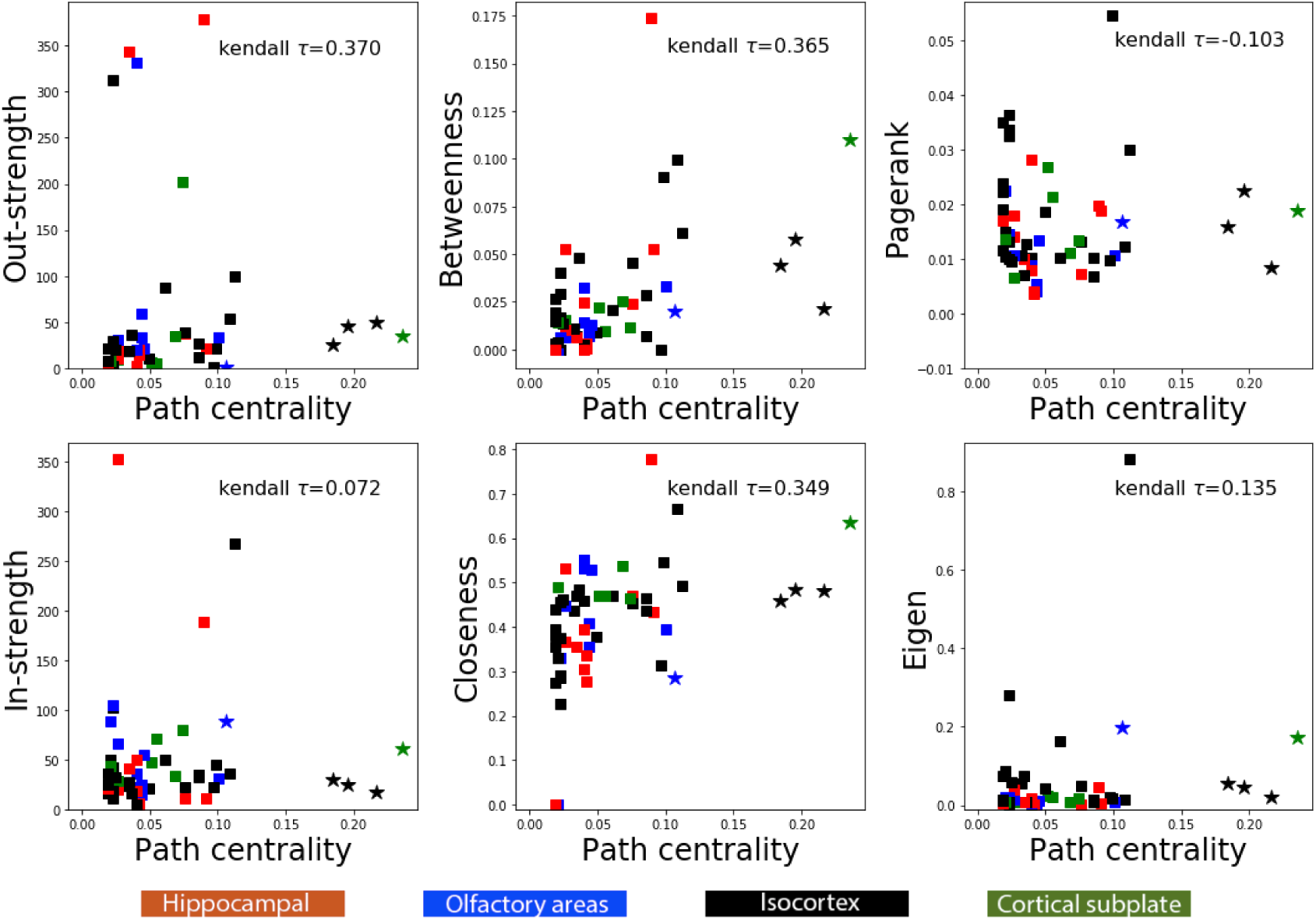
Path centrality compared to other centrality metrics. Each point corresponds to one of the 67 nodes in the network. The four star nodes constitute the hourglass core for *τ* =70%. All nodes are color-coded based on the broader brain region they belong to (isocortex, hippocampal formation, cortical subplate, olfactory areas).

## Footnotes

1 We use the terms “node”, “region” and ROI (“region of interest”) interchangeably.

2 We have repeated the analysis for other p-values in the range 0.01-0.1 – see Section SI.2.

3 The connection from AOB to COAp forms a single-edge bottleneck in the olfactory cascade. The weight of that connection in the Allen connectome is 0.46. With that value however the olfactory cascade require a different (lower) *θ* threshold than all other cascades. For this reason, we chose to artificially increase the weight of that connection from 0.46 to 1.

4 The probability for *l*_0_ = 0 is roughly 0.12 because the connectome has 616 connections and the average cascade includes only 75 of these edges – so the (unconditional) probability that a connection appears in at least one activation cascade is about 75/616*≈*0.12.

